# A Translational Porcine Model Linking Brain 5-HT_2A_ Receptor Occupancy to Psychedelic-like Behaviour

**DOI:** 10.64898/2026.11.16.767031

**Authors:** Clara A. Madsen, Miriam Navarro, Elisabeth Kolesnik, Julie F. M. Wedel-Heinen, Caroline Degel, Vanessa Drejfeldt, Virginia Camporesi, Despoina Kazani, Vladimir Shalgunov, Merlin Zabrocki, Matthias Herth, Dino Luethi, Friederike Holze, Matthias E. Liechti, Christoffer Bundgaard, Johan Juhl Weisser, Sys Stybe Johansen, Claus Svarer, Brice Ozenne, Hanne D. Garling, Gitte M. Knudsen

## Abstract

Many preclinical studies of serotonergic psychedelics use doses yielding drug exposures far exceeding the plasma concentrations associated with psychedelic effects in humans, limiting translational relevance. Here, we establish a translational pig model linking dose and plasma concentrations to in vivo and in vitro cerebral serotonin 2A (5-HT_2A_) receptor occupancy, enabling selection of pharmacologically relevant doses for porcine behavioural studies. Using [^11^C]Cimbi-36 positron emission tomography, we quantified brain 5-HT_2A_ receptor occupancy for five 5-HT_2A_ receptor agonists and two antagonists, finding good correspondence with human data. At doses achieving 5-HT_2A_ receptor occupancy tied to profound psychedelic experiences in humans, all five agonists induced headshakes in pigs, including lisuride and 2-Br-LSD, which are otherwise considered non-psychedelic agonists. These findings demonstrate that human-relevant 5-HT_2A_ receptor occupancy produces psychedelic-like behavioural effects in pigs, validating this model for translational evaluation and dosing of novel 5-HT_2A_ receptor compounds.

---

Serotonergic psychedelics, including psilocybin, lysergic acid diethylamide (LSD), and N,N-dimethyltryptamine (DMT), have regained scientific interest due to their potential for the treatment of neuropsychiatric disorders, often producing both immediate and long-lasting therapeutic effects after just one or two doses^1–4^. Stimulation of cerebral serotonin 2A (5-HT_2A_) receptors is a prerequisite for the acute psychedelic experience^5,6^, and the quality of the psychedelic experience has repeatedly been associated with the therapeutic outcome in a dose-dependent manner^7,8^. Recent positron emission tomography (PET) studies in humans have shown that the intensity of the subjective effects correlates with cerebral 5-HT_2A_ receptor occupancy and plasma drug concentration of LSD and psilocybin’s active metabolite, psilocin, and that 60–70% 5-HT_2A_ receptor occupancy is required for a profound psychedelic experience^9,10^, whereas higher doses involve more anxiety^11,12^. PET occupancy studies provide an opportunity not only to inform dose selection in clinical studies of novel psychedelics but also to guide preclinical studies toward translationally relevant drug doses and plasma concentrations.

While 5-HT_2A_ receptor stimulation does not cause motor behaviour in humans, it elicits a stereotypical behaviour in mice known as the head-twitch response (HTR), which is commonly used as a proxy for human psychedelic effects. HTR-like behaviour has been described in a range of different animals^13–18^, including pigs^19^. It is, however, unknown how the HTR relates to an animal’s subjective experience, and the HTR is not specific to 5-HT_2A_ receptor stimulation^13,20,21^. Preclinical studies have generated leading mechanistic hypotheses of psychedelic drug action, foremost that persistent therapeutic effects are caused by synaptic plasticity^22–24^. Yet, psychedelic drugs have demonstrated inconsistent therapeutic efficacy in animal models^25–33^, indicating that current preclinical approaches may not capture clinically relevant features. A major challenge is dose selection; preclinically, the drugs are often administered intravenously or intraperitoneally at doses much higher than those used clinically and chosen on the basis of how strongly they elicit the HTR which risks creating a translational gap. Moreover, appropriate dosing of novel 5-HT_2A_ receptor agonists associated with little or no psychedelic-like effects^33–35^ represents a separate challenge. Lisuride, an LSD analogue originally marketed for Parkinson’s disease, is widely labelled as a non-psychedelic 5-HT_2A_ receptor agonist, producing antidepressant-like effects in mice without eliciting HTR^30,32,36–39^. Another LSD analogue, 2-Br-LSD, produces no HTR but promotes synaptic plasticity in cultured neurons^40^ and prevents cluster headache attacks in humans without inducing psychedelic effects^41^. We therefore propose that plasma drug concentrations and in vivo target engagement of the cerebral 5-HT_2A_ receptor provide a more robust, species-independent foundation for comparisons of psychedelic and non-psychedelic compounds.

To this end, we determine the relationship between dose, plasma concentration, and in vivo cerebral 5-HT_2A_ receptor occupancy in pigs for three serotonergic psychedelics (DMT, psilocybin, and LSD), two putatively non-psychedelic compounds (lisuride and 2-Br-LSD), and two antagonists (ketanserin and volinanserin), using [^11^C]Cimbi-36 PET. We complement the in vivo measurements with in vitro competition binding against [^3^H]Cimbi-36 in porcine prefrontal cortex. We find that occupancy potencies and in vitro affinities fall within the range reported in humans, establishing the pig as a translational model for 5-HT_2A_ receptor pharmacology. Next, by administering doses achieving 5-HT_2A_ receptor occupancies associated with a profound psychedelic experience in humans, i.e., 60–70%, we show that all five agonists induce headshakes in pigs including, surprisingly, lisuride and 2-Br-LSD. The behavioural effects induced by 2-Br-LSD, but not lisuride, are reduced by volinanserin pretreatment, demonstrating their dependence on 5-HT_2A_ receptor activation. Together, these results support that in vivo target engagement, rather than stereotyped behaviour, can guide dose selection in preclinical psychedelic research.

## Results

### 5-HT2A receptor ligands dose-dependently occupy cerebral 5-HT_2A_ receptors in vivo in pigs

Thirty-six female pigs (weight 21 ± 2.8 kg (mean ± SD); approximately 8–10 weeks old) underwent [^11^C]Cimbi-36 PET scans to quantify cerebral 5-HT_2A_ receptor binding in terms of volume of distribution (V_T_). Each pig underwent in a single day both a baseline PET scan and one or two additional scans following intravenous drug administration. The number of scans, injected drug, and [^11^C]Cimbi-36 doses are given in Supplementary Table S1. One animal receiving a bolus of 1150 μg/kg DMT had a seizure-like reaction and was euthanised before the scan; scanner failure occurred for one animal receiving 30 μg/kg 2-Br-LSD; and two pigs were excluded from the analysis due to insufficient [^11^C]Cimbi-36 radiotracer production. When administered as a bolus (n_scans_ = 5), DMT did not generate occupancies >57% (Supplementary Figure S1), presumably due to its rapid elimination. Consequently, in the subsequent PET studies, DMT was administered as a bolus plus constant infusion, which generated steady-state plasma DMT levels (data not shown). For the analysis, 6 scans with DMT, 7 scans with psilocybin, 6 scans with LSD, 5 scans with lisuride, 6 scans with 2-Br-LSD, 5 scans with ketanserin and 5 scans with volinanserin were included.

We determined cerebral 5-HT_2A_ receptor occupancy by comparing [^11^C]Cimbi-36 PET V_T_ values at baseline and after drug administration, using the Lassen Plot^42^. We fitted the occupancies for each drug to the peak plasma drug concentration at the beginning of the scan (Figure 1C-D) or mean plasma drug concentration measured as an average during the entire scan (Supplementary Figure S2 and S3) to a Hill-Langmuir model. All drugs caused considerable dose-related 5-HT_2A_ receptor occupancies in the pig brain, and the relationship between peak or mean plasma drug concentrations and cerebral 5-HT_2A_ receptor occupancy conformed well to the Hill-Langmuir single-site binding model for all drugs. Individual occupancy values are shown in Supplementary Table S2. We also assessed the relationship between occupancy and drug doses and found good fits for both weight-adjusted and total doses (Supplementary Figure S2 and S3). We also estimated occupancies using a novel method that capitalises on shared data across studies, the Likelihood Estimation of Occupancy (LEO)^43^, which agreed closely with the Lassen plot estimates for our data (see Supplementary Table S2). For simplicity, we report the Lassen plot estimates throughout. Comparative half-maximal inhibitory plasma concentrations (IC_50_) and doses (ID_50_) in pigs and corresponding literature values in humans are shown in Table 1.

**Figure 1.**
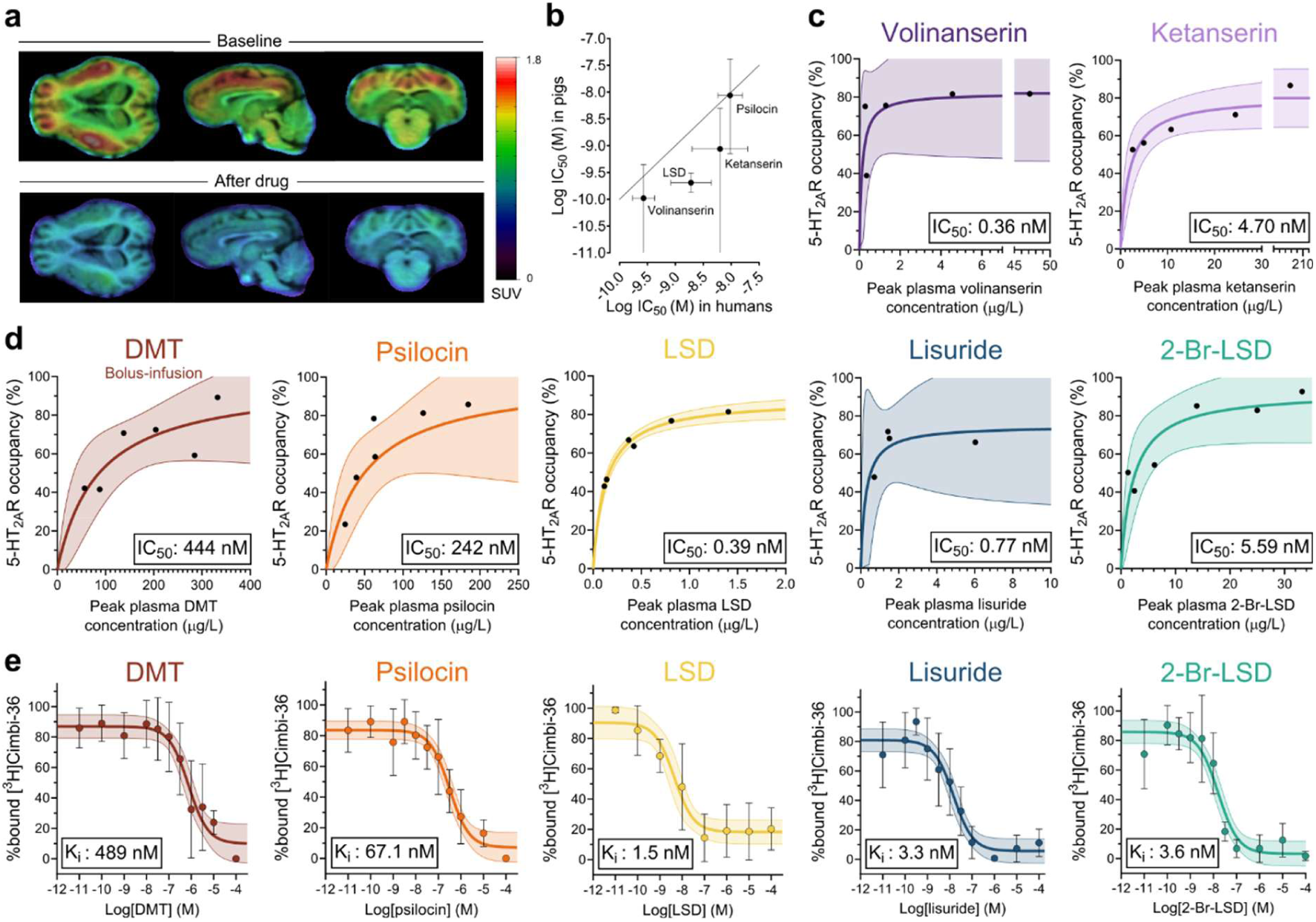
In vivo and in vitro 5-HT_2A_ receptor binding of DMT, psilocin, LSD, lisuride, 2-Br-LSD, volinanserin, and ketanserin in the pig brain. **a.** Representative SUV images of [^11^C]Cimbi-36 binding in the pig brain at baseline and after drug administration (100 μg/kg psilocybin) overlaid on a T1-weighted MRI image. The colour scale indicates SUV. **b.** Relationship between log IC_50_ (95% CI) in humans versus pigs for psilocin, LSD, ketanserin and volinanserin (see Table 1). Horizontal error bars show ± half the width of the published 95% CI on the log scale. For ketanserin and volinanserin, the lower confidence limit in pigs could not be determined; these bars extend to the axis limit. The solid line represents the line of identity. **c–d.** Relationship between peak plasma drug concentrations during the [^11^C]Cimbi-36 PET scan and cerebral 5-HT_2A_ receptor occupancy in the pig brain fitted with a Hill-Langmuir model. The solid lines show the model fits, and the 95% confidence interval bands are shown in coloured shading. **c.** Model fits for volinanserin revealed an IC_50_ (mean [95% CI]) of 0.13 [-0.11; 1.00] μg/L equivalent to 0.36 nM and a maximum occupancy (Occ_max_) (mean [95% CI]) of 82.2 [47.5; 122] %, and a goodness-of-fit of R^2^ = 0.39; and for ketanserin, an IC_50_ of 1.86 [0.23; 4.58] μg/L equivalent to 4.70 nM and an Occ_max_ of 80.6 [64.9; 98.5]%, and a goodness-of-fit of R^2^ = 0.83. **d.** Model fits for DMT revealed an IC_50_ of 83.7 [0.98; 688] μg/L equivalent to 444 nM and an Occ_max_ of 98.3 [59.3; 287]%, and a goodness-of-fit of R^2^ = 0.67; for psilocin, an IC_50_ of 49.4 [4.16; 79.4] μg/L equivalent to 242 nM and an Occ_max_ of 100% (constrained), and a goodness-of-fit of R^2^ = 0.74; for LSD, an IC_50_ of 0.13 [0.09; 0.17] μg/L equivalent to 0.39 nM and an Occ_max_ of 87.9 [81.6; 94.8]%, and a goodness-of-fit of R^2^ = 0.98; for lisuride, an IC_50_ of 0.26 [-0.32; 2.21] μg/L equivalent to 0.77 nM and an Occ_max_ of 75.0 [39.9; 136]%, and a goodness-of-fit of R^2^ = 0.50; for 2-Br-LSD, an IC_50_ of 2.25 [0.26; 9.75] μg/L equivalent to 5.59 nM and an Occ_max_ of 92.7 [67.0; 135]%, and a goodness-of-fit of R^2^ = 0.75. **e.** Comparable dose-dependent autoradiographic blocking of [^3^H]Cimbi-36 by DMT, psilocin, LSD, lisuride and 2-Br-LSD in pig prefrontal cortex. Data are presented as mean ± s.d.. The solid lines show the model fits, and the 95% confidence interval bands are shown in coloured shading.

**Table 1.**
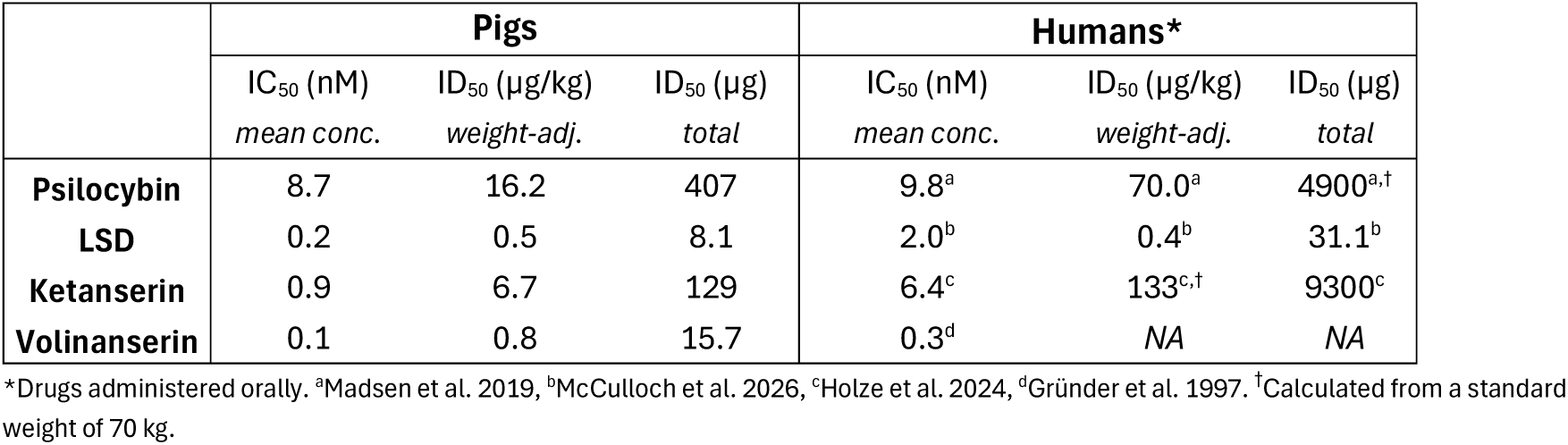
In vivo IC_50_ and ID_50_ of psilocybin, LSD, ketanserin, and volinanserin in pigs compared to literature data from human studies. IC_50_ values computed from mean plasma concentrations (*mean*) within the scan frame. ID_50_ values computed from weight-adjusted doses (*weight-adj.*) or total doses (*total*).

|  | **Pigs** |  |  | **Humans\*** |  |  |
|---|---|---|---|---|---|---|
|  | IC<sub>50</sub> (nM) | ID<sub>50</sub> (µg/kg) | ID<sub>50</sub> (µg) | IC<sub>50</sub> (nM) | ID<sub>50</sub> (µg/kg) | ID<sub>50</sub> (µg) |
|  | *mean conc.* | *weight-adj.* | *total* | *mean conc.* | *weight-adj.* | *total* |
| **Psilocybin** | 8.7 | 16.2 | 407 | 9.8<sup>a</sup> | 70.0<sup>a</sup> | 4900<sup>a,†</sup> |
| **LSD** | 0.2 | 0.5 | 8.1 | 2.0<sup>b</sup> | 0.4<sup>b</sup> | 31.1<sup>b</sup> |
| **Ketanserin** | 0.9 | 6.7 | 129 | 6.4<sup>c</sup> | 133<sup>c,†</sup> | 9300<sup>c</sup> |
| **Volinanserin** | 0.1 | 0.8 | 15.7 | 0.3<sup>d</sup> | *NA* | *NA* |

\*Drugs administered orally. <sup>a</sup>Madsen et al. 2019, <sup>b</sup>McCulloch et al. 2026, <sup>c</sup>Holze et al. 2024, <sup>d</sup>Gründer et al. 1997. <sup>†</sup>Calculated from a standard weight of 70 kg.

Arterial plasma samples acquired during PET scanning were used to assess plasma pharmacokinetics. Plasma drug concentrations peaked immediately after intravenous administration and the drugs were then eliminated with different half-lives (Table 2). All drugs displayed moderate inter-individual variability in elimination, and half-lives were dose-independent (Supplementary Table S2).

**Table 2.** In vivo plasma half-lives in pigs compared to literature data from human studies. Pig plasma half-lives (mean ± s.d.) were estimated from non-compartmental modelling.

|  | **Pigs** | **Humans** |
|---|---|---|
| **DMT** | 17.1 ± 4.4 min | 6 – 7 min<sup>a</sup> |
| **Psilocin** | 46.8 ± 17.1 min | 150 min<sup>b</sup> |
| **LSD** | 48.6 ± 13.4 min | 250 min<sup>b</sup> |
| **Lisuride** | 57.5 ± 12.6 min | - |
| **2-Br-LSD** | 59.7 ± 15.0 min | - |
| **Volinanserin** | 53.3 ± 5.4 min | 396 min<sup>c</sup> |
| **Ketanserin** | 38.2 ± 10.4 min | 210 min<sup>d</sup> |

<sup>a</sup> Erne et al. 2026, <sup>b</sup> Holze et al., 2022, <sup>c</sup> Gründer 1997, <sup>d</sup> Becker et al. 2023

### All 5-HT_2A_ receptor agonists acutely increase blood pressure in pigs

When given during the PET study, the agonists DMT, psilocybin, LSD, lisuride, and 2-Br-LSD all caused transient changes in blood pressure: DMT caused strong and immediate effects on blood pressure with an increase of up to 100%; psilocybin and LSD caused a moderate increase of up to 46% and 45%, respectively; lisuride and 2-Br-LSD were associated with small increases of up to only 20% and 21%, respectively (Supplementary Figure S4).

### In vitro affinity in pig brain tissue predicts in vivo potency

5-HT_2A_ receptor binding affinities for psychedelic drugs have been characterised by different laboratories in recombinant human receptor assays^44,45^ and rodent brain tissue^46–48^, but to relate in vivo potency and in vitro affinity in the same species and against the same radioligand, we performed competition binding against [^3^H]Cimbi-36 in pig prefrontal cortex. All assays were performed at physiological temperature, 37 °C, because assay temperature has previously been shown to affect binding affinities^49^.

All five agonists displayed complete blocking of [^3^H]Cimbi-36 binding and nanomolar affinities for the 5-HT_2A_ receptor (Figure 1E). The relationship between drug concentration and [^3^H]Cimbi-36 binding conformed well to a single-site binding model for all drugs. Assuming comparable plasma protein binding and extent of brain penetration, we found that in vitro affinities predicted the in vivo IC_50_ and potency rank order well: LSD > lisuride > 2-Br-LSD > psilocin > DMT, and the in vitro *K*_i_ values compare well with the in vivo IC_50_ values computed from peak plasma concentrations at scan start (Figure 1D-E), a proxy for the drug concentration the brain is exposed to in the beginning of the scan (mean IC_50_/*K*_i_ ratio 1.4; 95% limits of agreement 0.13–15, Supplementary Figure S6), with the differences lying within the uncertainty of the estimates. *K*_i_ values did not compare as well with IC_50_ values computed from mean plasma concentrations, and their differences deviated beyond measurement uncertainty (mean ratio IC_50_/*K*_i_ 0.39; 95% limits of agreement 0.013–12; Supplementary Figure S6). In vitro IC_50_ and *K*_i_ values for all drugs can be found in Supplementary Table S3.

### At matched 5-HT_2A_ receptor occupancy, both psychedelic and putative non-psychedelic compounds induce more headshakes than saline

Our group has previously shown that 80 μg/kg psilocybin (corresponding to an estimated 5-HT_2A_ receptor occupancy of 81%, see Supplementary Figure S2) elicits a stereotyped behavioural syndrome in pigs consisting of headshakes, scratching, and rubbing, lasting around 20 min^19^. On the basis of the in vivo occupancy data obtained in pigs in this study, we then assessed behaviours for DMT, psilocybin, LSD, lisuride, and 2-Br-LSD at doses corresponding to 60–70% 5-HT_2A_ receptor occupancy (see Supplementary Table S4), required to produce a profound psychedelic experience in humans^9,10^.

Nineteen female pigs (weight 19 ± 2.5 kg (mean ± SD); approximately 8–10 weeks old) were included in the behaviour study. Animals received the drugs in a randomized cross-over design: 400 μg/kg DMT, 30 µg/kg psilocybin, 0.6 µg/kg LSD, 5.8 µg/kg lisuride, and 11.7 µg/kg 2-Br-LSD. DMT was administered 30 min after monoamine oxidase inhibitor moclobemide (0.85 mg/kg), in an attempt to decrease the rate of elimination. We saw, however, no effect of moclobemide administration alone on the pigs’ behaviour (see Supplementary Figure S7) and the dose may have been insufficient to inhibit systemic degradation of DMT, especially because DMT with moclobemide had pharmacokinetics similar to that of DMT without moclobemide, as tested in a single pig (data not shown).

We found that within 60 min after intravenous administration to awake pigs, DMT (mean difference [95% CI]: 5.9 [3.5; 8.3], p = 0.002), psilocybin (mean difference [95% CI]: 6.9 [2.6; 11.2], p = 0.005), and LSD (mean difference [95% CI]: 10.4 [5.7; 15.2], p = 0.002) produced more headshakes than saline (Figure 2B). Interestingly, putative non-psychedelic drugs lisuride (mean difference [95% CI]: 30.2 [13.0; 47.5], p = 0.005) and 2-Br-LSD (mean difference [95% CI]: 3.6 [1.5; 5.6], p = 0.005) also produced more headshakes than saline within 60 min after administration (Figure 2B). When averaged across animals, the largest number of headshakes was seen after 4, 2, and 8 min for DMT, psilocybin, and LSD, respectively (Figure 2C). While for DMT, the number of headshakes was indistinguishable from baseline behaviour after 10 min, for psilocybin and LSD effects were more intermittent, lasting approximately 15 and 60 min, respectively. Only in the first 15 min after administration, all drugs induced significantly more headshakes than saline (all p-values <0.05) (Figure 2C). Unexpectedly, lisuride induced the highest number of headshakes, peaking 2 min after injection, while headshakes after 2-Br-LSD peaked after 4 min (Figure 2C), with effects returning to baseline levels after approximately 60 and 20 min, respectively. In fact, lisuride elicited more headshakes than saline within the 15–30 min (mean difference [95% CI]: 6.7 [1.9; 11.4], p = 0.03) and 30–45 min intervals (mean difference [95% CI]: 4.6 [1.6; 7.5], p = 0.02) (Figure 2C). At 45–60 min after administration, LSD (mean difference: 1.9, 95% CI [0.4; 3.4], p = 0.04) and lisuride (mean difference [95% CI]: 1.9 [0.4; 3.4], p = 0.04) elicited more headshakes than saline (Figure 2C).

**Figure 2.**
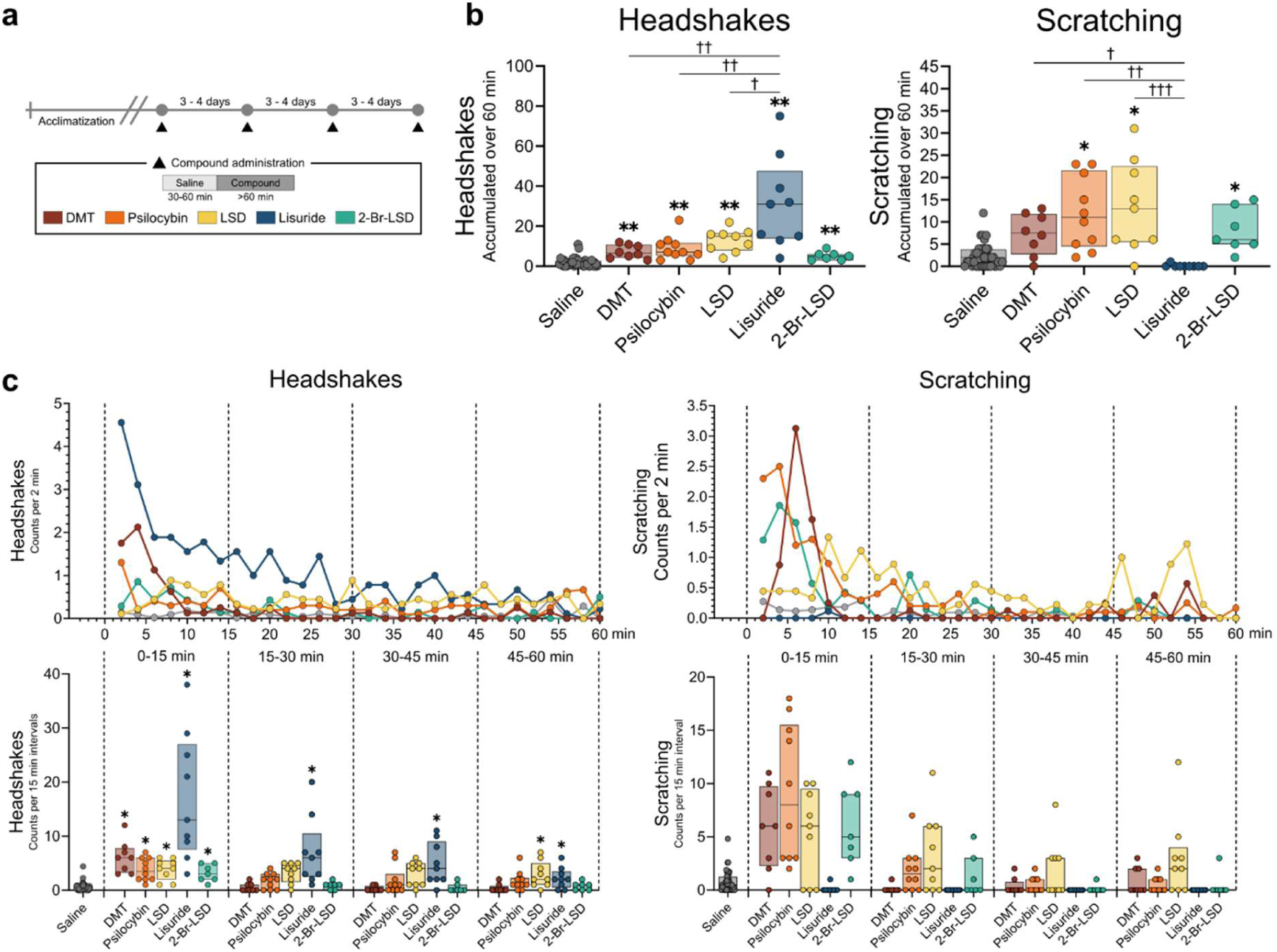
Pig behavioural effects after intravenous administration of DMT, psilocybin, LSD, 2-Br-LSD, and lisuride. **a.** Schematic showing the experimental setup and timeline for the drugs included in the analysis in this figure. **b.** Cumulative scores of headshakes and scratching during the first 60 min after administration compared to saline. **c.** Mean time-response curve of headshakes and scratching per 2 min during the first 60 min after administration, and cumulated scores of headshakes and scratching per 15 min intervals during the first 60 min after administration. Data are presented as median with 25th–75th percentiles. Drugs are compared with saline using paired t-tests, with significant differences marked with (*). Drugs are compared to each other using baseline-corrected linear mixed models, with significant ditferences marked with (†). ***/††† p<0.001, **/†† p<0.01, */† p<0.05.

Our linear mixed models revealed that scratching counts, but not headshakes, decreased across the experimental days for both animal cohorts (β =-0.7, s.e. = 0.3, 95% CI [-1.4; -0.1], p = 0.02) and (β =-0.6, s.e. = 0.3, 95% CI [-1.1; -0.02], p = 0.04), and the models were adjusted for this. Scratching behaviour was significantly increased relative to saline within 60 min after administration of psilocybin (mean difference [95% CI]: 9.4 [3.6; 15.2], p = 0.02), LSD (mean difference [95% CI]: 9.9 [1.8; 18.0], p = 0.04), and 2-Br-LSD (mean difference [95% CI]: 6.7 [2.7; 10.8], p = 0.02), but not lisuride or DMT (Figure 2B). We found that DMT (mean difference [95% CI]: 8.9 [0.1; 17.7], p = 0.048), psilocybin (mean difference [95% CI]: 13.0 [4.6; 21.3], p = 0.001) and LSD (mean difference [95% CI]: 15.9 [7.1; 24.7], p = 0.0002) all produced significantly more scratching than lisuride (Figure 2B). The largest number of scratching events, when averaged across animals, was seen after 6, 4, 10, and 4 min for DMT, psilocybin, LSD, and 2-Br-LSD, respectively, with effects returning to baseline levels after 10, 30, 60, and 30 min, respectively (Figure 2C). None of the drugs produced more scratching than saline in any of the 15-min intervals (all p-values >0.05) (Figure 2C). Rubbing occurred in some animals, but was not significantly difference from saline for any of the drugs (smallest p-value = 0.16, Supplementary Figure S5).

### The behavioural response to 2-Br-LSD, but not to lisuride, is reduced by volinanserin pretreatment

To test whether the behavioural effects induced by lisuride and 2-Br-LSD were caused by 5-HT_2A_ receptor stimulation, we blocked the receptor using the highly selective 5-HT_2A_ receptor antagonist volinanserin at a dose of 75 µg/kg. This dose was chosen based on the in vivo PET data with an estimated 5-HT_2A_ receptor occupancy of 85% (see Supplementary Figure S3) to ensure near-complete 5-HT_2A_ receptor blockade before administration of the agonists.

Volinanserin pretreatment caused a numerical reduction in the 2-Br-LSD-induced headshakes (Figure 3B and 3C), but we found no statistically significant difference between 2-Br-LSD with and without volinanserin pretreatment (mean difference [95% CI]: -3.1 [-13.8; 7.7], p = 0.7). However, 2-Br-LSD after volinanserin pretreatment was no longer significantly different to saline injection (mean difference [95% CI]: 1.4 [-1.1; 3.8], p = 0.2). Pretreatment also reduced 2-Br-LSD-induced scratching compared to no pretreatment (mean difference [95% CI]: -7.5 [-11.4; -3.6], p = 0.001), and 2-Br-LSD after volinanserin pretreatment was no longer significantly different from saline levels (2-Br-LSD+volinanserin vs. saline, p = 0.5). For lisuride, volinanserin pretreatment did not cause a statistically significant reduction in headshakes compared to no pretreatment (mean difference [95% CI]: 20.3 [-28.0; 68.7], p = 0.4), and lisuride with volinanserin pretreatment still induced significantly more headshakes than saline (mean difference [95% CI]: 25.5 [14.4; 36.6], p = 0.004) (Figure 3B). Lisuride did not induce scratching, neither before nor after volinanserin pretreatment (Figure 3B).

**Figure 3.**
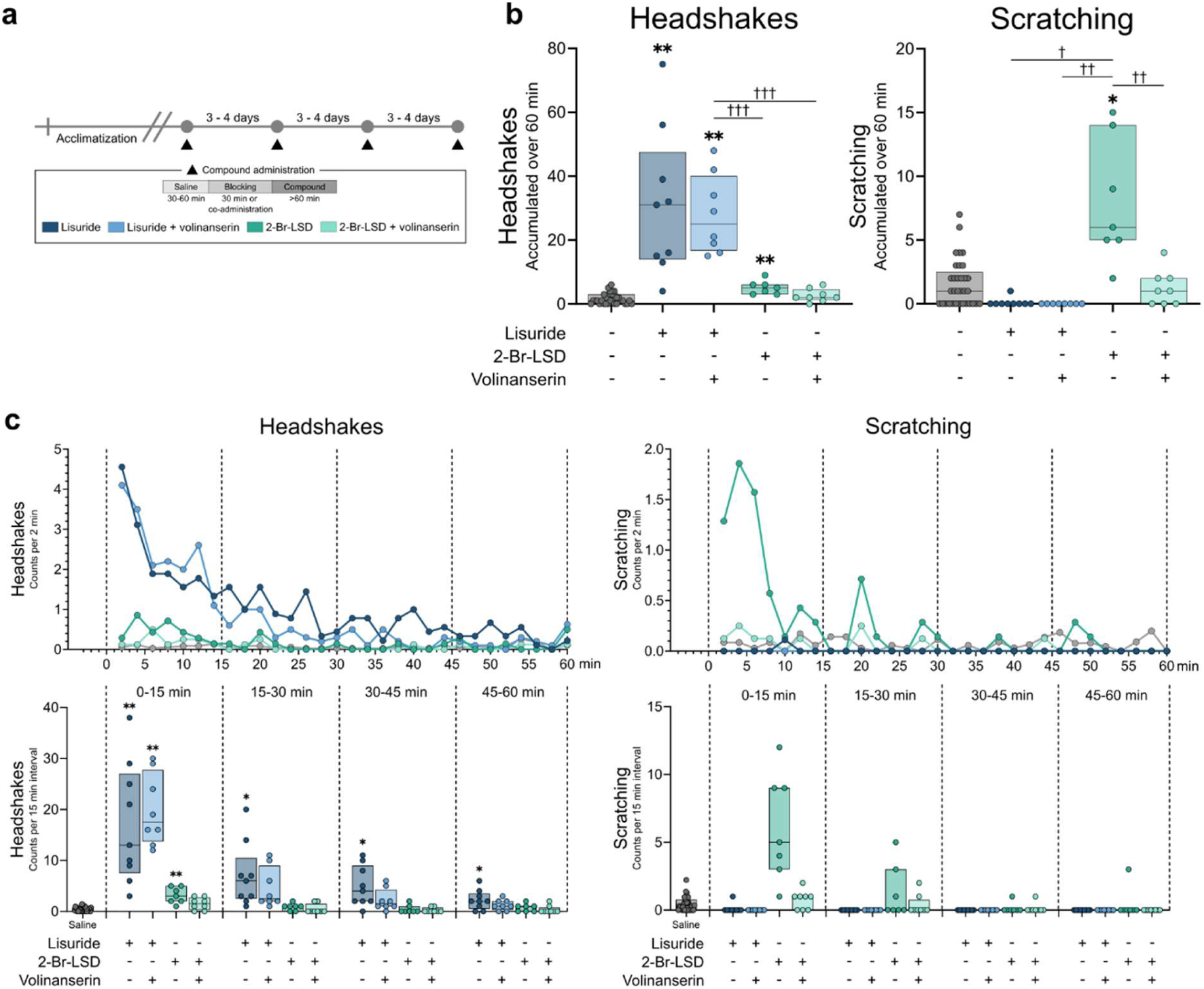
Pig behavioural effects after intravenous administration of 2-Br-LSD and lisuride before and after blocking with volinanserin. **a.** Schematics showing the experimental setup and timeline for the drugs included in the analysis in this figure. **b.** Cumulated scores of headshakes and scratching during the first 60 min after administration compared to saline. **c.** Mean time-response curve of headshakes and scratching per 2 min during the first 60 min after administration, and cumulated scores of headshakes and scratching per 15 min intervals during the first 60 min after administration. Data are presented as median with 25th–75th percentiles. Drugs are compared to saline using paired t-tests, with significant differences marked with (*). Drugs are compared to each other using baseline-corrected linear mixed models, with significant ditferences marked with (†). ***/††† p<0.001, **/†† p<0.01, */† p<0.05.

### LSD has slow pharmacokinetics and pharmacodynamics in pigs

To assess time-dependent effects of DMT, lisuride and LSD on receptor occupancy, we performed follow-up scans on the same day after plasma drug concentrations had decreased in one animal receiving a DMT bolus-infusion, two animals receiving LSD, and two animals receiving lisuride. Whereas follow-up scan occupancies for DMT and lisuride fell within the 95% prediction band of the relationship established from the first block scans (Figure 4A), LSD follow-up scans showed higher occupancies than predicted from the concomitant plasma concentrations, i.e., the occupancy remained high after plasma concentrations had decreased substantially (Figure 4A).

**Figure 4.**
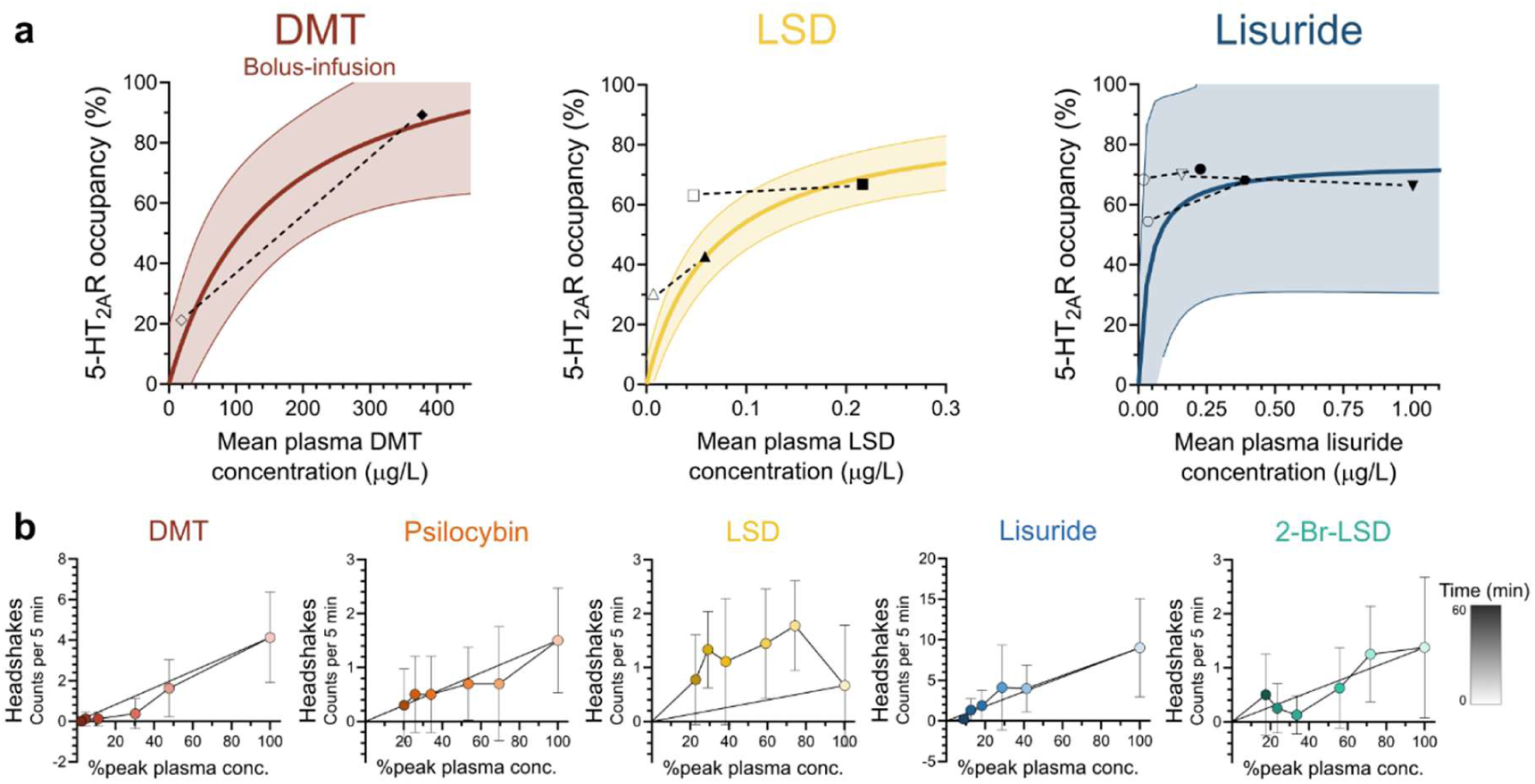
Relationship between pharmacokinetics and pharmacodynamics in pigs. **a.** 5-HT_2A_ receptor occupancy from follow-up scans for DMT, LSD, and lisuride. Relationship between mean plasma drug concentrations over the 90 min [^11^C]Cimbi-36 PET scan and cerebral 5-HT_2A_ receptor occupancy. Model fits and prediction bands represent the data from the first block scan. Filled symbols represent the occupancy measured in the first scan, while the connected empty symbols represent the occupancy measured in the follow-up scan in the same animal. **b.** Relationship between plasma concentrations and headshakes at 5, 10, 15, 30, 45, and 60 min after drug administration. Data points are chronologically connected by black lines, and points are coloured by time, i.e., the number of minutes post-drug administration. Percentage peak plasma concentrations are extrapolated from normalised concentration values from PET experiments fitted to a two-phase decay model (see Supplementary Figure S9).

We also assessed the relationship between plasma concentrations and behavioural effects over time. For LSD, we found an anticlockwise hysteresis loop, in which psychedelic-like effects remained high while plasma concentrations decreased (Figure 4B), consistent with observations in humans^10,50^. This effect was not observed for any of the other compounds, which all showed a direct relationship between plasma concentrations and psychedelic-like effects, independent of time after dosing.

## Discussion

In our pharmacological characterization of seven 5-HT_2A_ receptor ligands, we report the relationship between drug dose, plasma concentrations, and in vivo cerebral 5-HT_2A_ receptor occupancy of three serotonergic psychedelics (DMT, psilocybin, and LSD), two putatively non-psychedelic compounds (lisuride and 2-Br-LSD), and two 5-HT_2A_ receptor antagonists (ketanserin and volinanserin) in pigs. At doses achieving 5-HT_2A_ receptor occupancy associated with profound psychedelic experiences in humans, all five agonists induced headshakes, including lisuride and 2-Br-LSD, which until now have been considered non-psychedelic because they do not elicit HTR in mice^32,36–40^.

Our in vivo IC_50_ estimates in pigs were within one order of magnitude of previously reported human in vivo estimates for psilocin^9^, LSD^10^, ketanserin^51^ and volinanserin^52^ (see Figure 1B and Table 1), supporting the translational relevance of the pig model for quantifying in vivo cerebral 5-HT_2A_ receptor target engagement. The in vivo IC_50_ estimates in pigs were generally lower than those in humans, consistent with the way IC_50_ values were determined, namely on the basis of average plasma drug concentration over the course of the PET scan. The different routes of administration (intravenous in pigs vs oral in humans), and faster drug metabolism in pigs than in humans cause a lower accumulated drug exposure to the pig brain which likely lead to an underestimation of IC_50_ values in pigs. Similarly, our in vitro *K*_i_ values agree with those from cell studies using the recombinant human 5-HT_2A_ receptor for DMT, psilocin, LSD^44,45^, lisuride^53,54^, and 2-Br-LSD^40^, indicating that porcine and human 5-HT_2A_ receptor pharmacology is very similar. This observation is consistent with the amino acid sequence for the 5-HT_2A_ receptor being highly conserved between humans and pigs^55^. By contrast, mice and rats carry a serine-to-alanine substitution within the receptor (S242A), at a site important for ergoline and tryptamine ligand binding^56^; this substitution is not present in the porcine or human receptor^55^. We also find good agreement between in vitro *K*_i_ values and in vivo IC_50_ values, with a preserved rank order across the five agonists. Interestingly, binding affinities from cell studies measured at the recombinant human 5-HT_2A_ receptor have previously been shown to correlate with in vivo estimated average doses at which the drugs are psychoactive in humans^45,57^. However, while in vitro studies are performed under controlled equilibrium binding conditions, in vivo studies are done under non-equilibrium conditions and affected by factors such as protein binding in blood and brain interstitial fluid and non-displaceable binding. Thus, although in vitro affinity may provide a useful indication of relative in vivo potency, direct measurement of cerebral receptor occupancy remains essential for establishing in vivo target engagement.

Although DMT, psilocybin, and LSD induce comparable levels of headshakes in pigs at 60–70% cerebral 5-HT_2A_ receptor occupancy, the effects of DMT and psilocybin resolved faster than after LSD. These findings mirror human data indicating that at 60–70% 5-HT_2A_ receptor occupancy (corresponding to oral doses of up to 30 mg psilocybin or 100 µg LSD^9,10^), both drugs induce similar subjective intensities, but with a longer duration for LSD^12^. Intriguingly, at doses matched on the basis of 5-HT_2A_ receptor occupancy, we find that the putative non-psychedelic compounds 2-Br-LSD and lisuride cause a behavioural syndrome in pigs similar to that of the serotonergic psychedelics. As reviewed in Kehler and Lindskov, 2025, non-controlled humans studies demonstrate that a range of lisuride doses produce psychedelic-like effects^58^. For 2-Br-LSD, a single oral dose of 500 μg induced an “LSD-like delirium” lasting at least seven hours^59^, while lower doses (30 µg/kg) caused only “flabby” or “light drunk” sensations^41^. This dose-dependent variation suggests that these analogues can induce psychedelic effects in humans, provided they are administered at doses that achieve relevant cerebral 5-HT_2A_ receptor occupancy. By contrast, neither lisuride^60–62^ nor 2-Br-LSD^40^ induce HTR in mice, at least not at the doses tested. In the absence of HTR, lisuride still induces other behavioural and EEG effects, associated with psychoactivity^62^, demonstrating the limitations of the HTR. Although on logarithmic scales, HTR potency in mice correlates with self-reported recreational doses of psychedelics in humans^63^, there are exceptions. For instance, fenfluramine^20^ and 5-hydroxytryptophan^13,21^ elicits HTR in rodents despite being non-psychedelic in humans. Further, HTR in mice may be insufficient to determine whether a compound is psychedelic because a drug may act on multiple receptors that collectively contribute to the interoceptive stimulus^64^. Our data support this notion: While we find that the highly specific 5-HT_2A_ receptor antagonist volinanserin blocks 2-Br-LSD-induced headshakes and scratching, suggesting that these are mediated by 5-HT_2A_ receptor agonism, lisuride-induced headshakes were unaltered suggesting that lisuride-induced headshakes are mediated by non-5-HT_2A_ receptor mechanisms. Since lisuride has higher affinity for 5-HT_1A_ and dopamine 2 (D_2_) receptors than for 5-HT_2A_ receptors^35,54^, doses generating 60–70% 5-HT_2A_ receptor occupancy are likely to engage with these other targets in vivo. Both receptors have previously been shown to be involved in the rodent HTR; 5-HT_1A_ receptor antagonism has been shown to increase the HTR to both LSD and (-)-2,5-dimethoxy-4-methylamphetamine (DOM), whereas 5-HT_1A_ receptor agonism decreases it^53,65^. Accordingly, lisuride’s 5-HT_1A_ receptor stimulation has been suggested to explain why it does not induce HTR in mice^53^. In addition, some non-specific D_2_ antagonists have been shown to block 2,5-dimethoxy-4-iodoamphetamine (DOI)-induced HTR in rodents^14,66^, suggesting that this receptor could also contribute to this behaviour. We also find that psilocybin, LSD, and 2-Br-LSD all induced scratching behaviour, while lisuride did not. A similar behavioural phenomenon, ear scratching, is seen in mice after administration of certain phenethylamine-type psychedelics but not after tryptamines, because 5-HT_1B_ receptor agonism blocks this response in mice^44,67^. For this reason, ear scratching has been used less consistently than the HTR in the literature. Similar to lisuride, we have previously shown that another putative non-psychedelic compound, tabernanthalog, produces headshakes but not scratching in pigs at a dose estimated to generate 45% 5-HT_2A_ receptor occupancy^68^. Together, our findings highlight that in vivo cerebral 5-HT_2A_ receptor engagement, eventually combined with a behavioural assay in a large animal model are likely to provide a more reliable basis for dose selection and for the evaluation of putative non-psychedelic compounds in humans.

Our pharmacokinetic investigation revealed that drug half-lives in the anesthetised pigs generally are about 3–7 times shorter than those reported in humans, except for DMT, which has a half-life that is about 2.6 times longer in pigs (see Table 2). Species differences in drug half-lives are common: Mice metabolise psilocin at twice the rate of humans^69^, and in nonhuman primates, psilocin has a half-life of 49 min^70^, similar to what we observe in pigs. We also demonstrate that the temporal course of the porcine behavioural responses also corresponds to what has been reported for the human psychedelic experience. Following intravenous administration in humans, 5–25 mg DMT produces subjective effects almost immediately, which peak at approximately 2 min and subside completely within 12–30 min^71^. Similarly, intravenous administration of 1.5–2 mg psilocybin produces subjective effects almost immediately, peaking at approximately 4 min and subsiding completely within 20 min^72^. In contrast, intravenous administration of 80 µg LSD (free-base equivalent) produces subjective effects that peak at approximately 1.2 h and subside within 8 h^73^, although a delay of approximately 10 min in the onset of subjective effects has also been reported^74^. Human studies show that LSD exhibits not only slow pharmacokinetics and pharmacodynamics but also a temporal dissociation between the two, with a lag between plasma concentrations and both 5-HT_2A_ receptor occupancy^10^ and subjective effects^10,50^. This temporal lag has not been observed with DMT^71^ or psilocybin^10^. Here, we show that the relationship between plasma concentrations and headshakes in pigs likewise exhibits clear anticlockwise hysteresis for LSD, but not for any of the other compounds. Furthermore, we demonstrate that high 5-HT_2A_ receptor occupancy is maintained despite a substantial decline in plasma LSD concentrations, a dissociation that parallels the observed behavioural hysteresis. LSD was also the only compound to produce a delayed effect on blood pressure. Assuming that the effects are mediated, at least in part, by direct vasoconstriction resulting from 5-HT_2A_ receptor stimulation on blood vessels^75,76^, this finding argues against slow blood-brain barrier penetration as the primary explanation for LSD’s temporal lag and instead supports a role for slow binding kinetics of LSD at the receptor. Consistent with this interpretation, in vitro studies have demonstrated unusually slow binding kinetics of LSD at the 5-HT_2A_ receptor, including a slow dissociation rate that prolongs receptor residence time. Structural and mutagenesis studies further implicate specific residues in the receptor’s second extracellular loop as key determinants of the slow off-rate^77,78^.

This study has several strengths. We used state-of-the-art neuroimaging methodology by conducting PET imaging to estimate receptor occupancy using the well-validated agonist 5-HT_2A_ receptor radioligand [^11^C]Cimbi-36, which is currently the only agonist radiotracer available for neuroimaging of the 5-HT_2A_ receptor. The PET imaging was conducted in a similar manner to previous human studies^9,10,51^, which allows for direct cross-species comparison, whereas the only other preclinical studies that have investigated 5-HT_2A_ receptor occupancy of psychedelics to date rely on antagonist radiotracers or ex vivo methods^48,79–81^. Use of the pig as the experimental model is an additional strength, because the pig brain is gyrencephalic and closer to the human brain. The porcine model also enables repeated intravenous dosing, arterial blood sampling, same-day scanning, and post-mortem tissue collection in the same animal. This combination of experimental procedures is difficult to achieve in nonhuman primates and generally not feasible in rodents, given the blood volumes required for pharmacokinetic characterisation.

There are also limitations to the study. First, behavioural assessments were performed following intravenous bolus administration of DMT, whereas 5-HT_2A_ receptor occupancy at the behavioural dose was estimated from the plasma concentration-occupancy relationship established during bolus-infusion PET experiments. Infusion of DMT in awake pigs was not possible because it interfered with the behavioural outcomes. Second, plasma pharmacokinetics were characterised under anaesthesia, which may have altered drug metabolism and, consequently, may not fully reflect pharmacokinetics in awake animals or humans. Finally, the behavioural experiments employed a randomised cross-over design, meaning that carryover effects from previous drug administrations cannot be excluded. However, sensitivity analyses provided no evidence of a substantial effects of the day of administration on headshakes.

In conclusion, our study is the first to characterise the relationship between drug dose, plasma concentration, and in vivo cerebral 5-HT_2A_ receptor occupancy of three serotonergic psychedelics (DMT, psilocybin, and LSD), two putatively non-psychedelic compounds (lisuride and 2-Br-LSD), and two 5-HT_2A_ receptor antagonists (ketanserin and volinanserin) in a highly translational pig model. Compared with the other compounds tested, LSD exhibited slower pharmacokinetics and pharmacodynamics with a temporal lag between plasma concentrations and drug effects. Using in vivo IC_50_ values derived from the PET study to select doses for characterising acute behavioural effects, we found that all five agonists induced headshakes when administered at doses matched for 5-HT_2A_ receptor occupancy. Contrary to previous findings, our results indicate that 2-Br-LSD closely resembles the other psychedelics in pigs, inducing both headshakes and scratching, which were blocked by pretreatment with volinanserin. In contrast, lisuride induced a rapid and pronounced increase in headshakes, exceeding that observed with any of the other agonists, but did not induce scratching; moreover, the headshake response was not blocked by volinanserin. Notably, the pharmacokinetic, pharmacodynamic, and behavioural profiles of DMT, psilocybin, and LSD in pigs closely mirrored those reported in humans. Overall, our findings establish the pig as a valuable translational model for investigating the pharmacokinetics, pharmacodynamics, and behavioural effects of serotonergic compounds, and highlight the importance of integrating in vivo cerebral 5-HT_2A_ receptor occupancy with behavioural assessments to improve dose selection and interpretation of preclinical findings in psychedelic research.

## Methods

### Animals

Female domestic pigs (crossbreed of Landrace × Yorkshire × Duroc) were obtained from a local farm and acclimatised for at least one week in an enriched environment with a 12-h light/dark cycle, free access to water, and food twice daily. Procedures were approved by the Danish Council of Animal Ethics (Journal no. 2022-15-0201-01156), performed in accordance with European Commission Directive 2010/63/EU, and are reported in compliance with the ARRIVE guidelines.

### Drugs

Psilocybin (produced by University of Chemistry and Technology and the National Institute of Mental Health, Prague, Czech Republic), LSD (1 mg/mL in acetonitrile for in vivo experiments; Lipomed, LGC GmbH, Luckenwalde, Germany, or LSD freebase for in vitro experiments; kindly supplied by Terran Biosciences Inc., Miami, USA), DMT (kindly supplied by Terran Biosciences Inc., Miami, USA), 2-Br-LSD (Toronto Research Chemicals, Vaughan, Canada), lisuride maleate (Tocris Bioscience, Bristol, United Kingdom), ketanserin tartrate (Sigma-Aldrich, Saint Louis, USA), volinanserin (MDL 100,907 hydrochloride; Hello Bio, Dunshaughlin, Ireland), and moclobemide (Acros Organics BV, Geel, Belgium) were dissolved in ≤5% DMSO (except LSD) and diluted with 0.9% saline to 10–20 mL before injection.

### PET scanning

Each pig underwent two to three [^11^C]Cimbi-36 PET scans on the same day: a baseline scan followed by one or two scans preceded by drug administration. The drugs were administered as intravenous bolus injections through one of the mammary veins over approximately 30–60 s. We used [^11^C]Cimbi-36 in preference to antagonist radiotracers such as [^11^C]MDL100907 or [^18^F]altanserin because agonist radioligands preferentially label the high-affinity, G protein-coupled state of the receptor and therefore provide a more functionally meaningful estimate of the receptor pool available to agonist drugs^82,83^. Drug administration timing relative to tracer injection was determined from each drug’s expected pharmacokinetics: psilocybin 30 s, LSD 25 min, lisuride 5 min, 2-Br-LSD 5 min, ketanserin 10 min and volinanserin 10 min before tracer injection. DMT was given either as a bolus 30 s before tracer injection or as a bolus plus constant infusion beginning 5–15 min before. For two LSD, one DMT bolus-plus-infusion, and three lisuride administrations, a follow-up scan was acquired 1 h after the end of the first blocking scan to obtain two occupancy measurements from a single administration. Drug doses are given in Supplementary Table S1.

#### Animal preparation

Animals were prepared as previously described^19^. Briefly, anaesthesia was induced with 0.14 mg/kg intramuscular Zoletil mixture (11.36 mg/mL xylazine, 11.36 mg/mL ketamine, 1.82 mg/mL butorphanol, 1.82 mg/mL methadone) and maintained with intravenous propofol (12.5–15.0 mg/kg/h) and fentanyl (5 µg/kg/h). Animals were endotracheally intubated and ventilated with ∼34% oxygen in normal air at 10–12 mL/kg. Arterial access was established in both femoral arteries (Teleflex, Wayne, PA, USA) and venous access in both mammary veins (neonatal feeding tube; Vygon Denmark, Copenhagen, Denmark) and in the ear veins. A urinary catheter was placed to avoid discomfort. Vital parameters including heart rate, blood pressure, O_2_ saturation, end-tidal CO_2_, body temperature, and blood glucose levels, were monitored throughout the PET scanning. Immediately after scanning, animals were euthanised by intravenous injection of 15–20 mL pentobarbital (400 mg/mL, ScanVet Animal Health, Fredensborg, Denmark) and the brains were immediately extracted, separated into hemispheres, snap-frozen on powdered dry ice, and stored at −80 °C.

#### Scanning protocol and radiochemistry

Pigs were scanned in a Siemens high-resolution research tomograph (HRRT; Siemens, Malvern, PA, USA). [^11^C]Cimbi-36 was prepared at the Department of Clinical Physiology and Nuclear Medicine, Copenhagen University Hospital Rigshospitalet, as previously described^84^, using carbon-11 methyl triflate, and injected intravenously as a bolus over approximately 20 s through one of the mammary veins. The PET acquisition was started simultaneously with injection of the radiotracer, and the pigs were scanned for 90 min.

#### Arterial input function

Arterial blood radioactivity was measured continuously every 2000 ms for the first 2000 s using an ABSS autosampler (Allogg Technology, Strängnäs, Sweden), and manual heparinised samples were drawn at 2.5, 5, 10, 20, 30, 40, 50, 70, and 90 min. Whole-blood and plasma radioactivity were counted in an automated gamma counter (Cobra 5003, Packard Instruments; or Hidex AMG, Hidex Oy, Turku, Finland) cross-calibrated to the HRRT. Radiolabelled parent and metabolite fractions were determined in plasma using radio-high-performance liquid chromatography (HPLC) as previously described^85^. Briefly, plasma was filtered through a 0.45 µm syringe filter (Whatman GD/X 13 mm, Cytiva) and subsequently diluted 1:1 with 20 mM phosphate buffer and 5 mM sodium-1-decanesulfonate pH 7.2 with 2% isopropanol. Samples were analysed in a fully automated column-switching HPLC system (UltiMate 3000, Thermo Fisher Scientific) connected to a radio-HPLC detector (PosiRam Model 4, LabLogic Systems). The HPLC system was equipped with a small extraction column (Shimpack MAYI-ODS 30×4.6 mm, Shimadzu Corporation) combined with an analytical column (Onyx Monolithic C18 50×4.6 mm, Phenomenex). The extraction mobile phase consisted of 100% phosphate buffer (composition mentioned above), while the elution mobile phase consisted of 71% phosphate buffer (100 mM) and 2 mM sodium 1-decanesulfonate pH 2.6 and 29% acetonitrile. Samples were injected in a volume of 4 mL, and the analysis was run at a flow of 5 mL/min at 25 °C. Total runtime for each sample was 8.55 min with a 4-min extraction step, a 4-min elution step, and 0.55 min of equilibration. Four eluate fractions were collected from the 20-min timepoint at 2 min intervals using a fraction collector (Foxy Jr FC144; Teledyne) and radioactivity subsequently measured using a gamma counter (Wizard 2480, Perkin Elmer). The parent fraction was calculated as the radioactivity of the parent eluate divided by the total collected radioactivity.

#### Image reconstruction and kinetic modelling

List-mode data were reconstructed into 38 frames (6×10, 6×20, 4×30, 9×60, 2×180, 8×300, 3×600 s) using OP-3D-OSEM with point spread function modelling (16 subsets, 10 iterations) and standard corrections^86^, with attenuation correction by MAP-TR µ-map^87^. Each time frame comprised 207 planes of 256 × 256 voxels of 1.22 × 1.22 × 1.22 mm. Brain parcellation was done using a previously published automatic pig brain delineation method for PET images^88^. Kinetic modelling was performed using the kinfitr R package (v. 0.9.1)^89,90^. Total volume of distribution (V_T_) was estimated by Logan graphical analysis^91^ using a metabolite-corrected arterial input function, with a T* of 9 frames (corresponding to 30–90 min). 5-HT_2A_ receptor occupancy was calculated using the Lassen plot^42^ using regional V_T_ values from thalamus, dorsal striatum, somatosensory cortex, frontal cortex, insular cortex, occipital cortex, temporal cortex, cingulate cortex, hippocampus, white matter, and cerebellum (excluding the vermis). We also estimated occupancies using a novel method, Likelihood Estimation of Occupancy (LEO), where non-displaceable distribution volume (V_ND_) and occupancy are estimated using direct maximum likelihood estimation^43^. LEO can provide more accurate and precise estimates of occupancy and V_ND_ because it uses a covariance matrix derived, for example, from a test-retest dataset. We used a covariance matrix derived from eight test-retest [^11^C]Cimbi-36 datasets acquired as above.

#### Occupancy-concentration modelling

Peak plasma drug concentrations were defined as the highest plasma concentration after scan start. Peak plasma drug concentrations for DMT (bolus-infusion), LSD, lisuride, 2-Br-LSD, volinanserin, and ketanserin were measured in arterial blood samples taken immediately prior to scan start. For psilocin and DMT (bolus), because the scan was started immediately after drug injection, continuous arterial sampling was performed for the first 3 min after drug injection to give an average drug concentration at this time. Mean plasma drug concentrations were calculated as the area under the plasma concentration-time curve within the scan timeframe, divided by scan duration (90 min). The relationship between 5-HT_2A_ receptor occupancy and plasma concentrations was modelled using the Hill-Langmuir model:

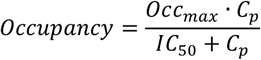

where Occ_max_ is the theoretical maximal occupancy, C_p_ is the plasma drug concentration, and IC_50_ the plasma concentration producing half-maximal occupancy. Curve fitting was performed in GraphPad Prism (v11.0.2; GraphPad Software, San Diego, CA, USA). Occ_max_ was bounded between 0 and 100%, with unbounded 95% confidence intervals (CI).

### Plasma drug concentrations

Plasma exposures were determined by extracting analytes from arterial plasma using protein precipitation followed by liquid chromatography–tandem mass spectrometry analysis. Mass spectrometry quantification was performed using multiple reaction monitoring in positive ionization mode. For DMT, psilocin, and LSD, heavy-isotope-labelled analogues were used as internal standards (DMT-d_6_, psilocin-d_10_, LSD-d_3_). The internal standards were LSD-d3 for 2-Br-LSD, trans-dihydro lisuride-d_10_ for lisuride, risperidone for ketanserin, and an internal Lundbeck compound for volinanserin. Methanol was used as precipitation solvent for DMT, LSD and 2-Br-LSD, whereas acetonitrile was used for psilocin, lisuride, ketanserin and volinanserin. The measurement range were: 0.25–250 ng/mL for DMT; 0.102-204 ng/mL for psilocin; 0.01-10 ng/mL for LSD; 0.05-10 ng/mL for 2-Br-LSD; 0.01–10 ng/mL for lisuride; 0.1–1000 ng/mL for ketanserin and 0.01–10 ng/mL for volinanserin. Methodologies for determination of DMT^92^, psilocin^9^, LSD^93^ and ketanserin^6^ in plasma have previously been described in the literature. The methodology for analysis of 2-Br-LSD was adapted from the LSD methodology^93^. Lisuride and volinanserin were analysed by adapting a protocol for the analysis of ivabradine^94^.

Pharmacokinetic parameters, including half-lives and area under curve (AUC), were estimated by non-compartmental analysis using the PKNCA R Package (v. 0.12.1)^95^.

### Cardiovascular effects

Heart rate (HR), systolic blood pressure (SBP), diastolic blood pressure (DBP), and mean arterial pressure (MAP) were recorded throughout PET scanning. For each animal, baseline was calculated as the mean of all pre-injection measurements within 60 min before drug administration; where no measurement fell in that window, the two measurements preceding administration were used. Percent change from baseline was calculated as:

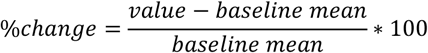

### Radioligand binding assay

Prefrontal cortex from five pig brains that had undergone PET scanning without drug intervention was sectioned at 20 µm on a cryostat (CryoStar NX50, Thermo Fisher Scientific) at -12 °C with a blade temperature of -20 °C, thaw-mounted onto a glass slide (StarFrost® Adhesive Slides with silane coating, Walder Knittel Glasbearbeitungs GmbH, Braunschweig, Germany), air-dried, and stored at -20 °C until use. Sections were thawed for 30–60 min, pre-incubated twice for 10 min in assay buffer (50 mM Tris-HCl, 4 mM CaCl2, 0.1% BSA, 0.01% ascorbic acid, pH 7.4) at 37 °C, then incubated with 1 nM [^3^H]Cimbi-36 (Novandi Chemistry, Södertälje, Sweden) and increasing concentrations (10−¹¹–10−⁴ M) of DMT, psilocin, LSD, 2-Br-LSD, or lisuride for 30 min at 37 °C on an orbital shaker. Incubation was terminated by two 10-min washes in ice-cold 50 mM Tris-HCl and a rapid rinse in ice-cold deionised water. The slides were air-dried for a minimum of 45 min and fixed overnight in a paraformaldehyde vapour chamber at 4 °C. The next day, the samples were moved to an desiccator for 60 min to remove any excess moisture and then placed in a radiation-shielded image plate cassette (Fuji Cassette2 BAS-TR 2040, Fujifilm, Tokyo, Japan) along with two radioactive tritium standards ([^3^H]microscales ART0123B (3.0–109.4 nCi/mg) and ART0123C (0.10-15.9 nCi/mg), American Radiolabeled Chemicals, Inc., St. Louis, USA). The slides were exposed to a tritium-sensitive image plate (Fuji Imaging Plate BAS-TR, Fujifilm, Tokyo, Japan) for 7 days at 4 °C. The imaging plate was scanned on an Amersham™ Typhoon™ IP Biomolecular Imager (GE healthcare, Chicago, USA). Regions of interest were delineated in ImageJ (NIH, Bethesda, MD, USA) and grey values were converted to nCi/mg tissue equivalent using a four-parameter fit (Rodbard NIH) of the decay-corrected tritium standards, then to fmol/mg tissue equivalent using the decay-corrected molar activity. Total binding was measured in cortical grey matter and non-specific binding in white matter of the same section. Specific binding was calculated as the difference between total binding and non-specific binding. GraphPad Prism (v11.0.2) was used to calculate IC_50_ values using a one-site competition model:

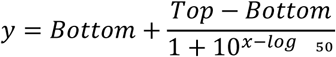

Corresponding inhibitory constant (K_i_) values were calculated using the Cheng-Prusoff equation:

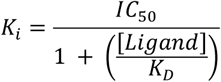

K_D_ for [^3^H]Cimbi-36 was determined in a saturation binding experiment and was set to 0.26 nM (data not shown). For visualisations, specific binding was normalised per animal so that the lowest and highest values in the dataset corresponded to 0% and 100%.

Agreement between in vitro *K*_i_ and in vivo IC_50_ values was assessed by Bland–Altman analysis of log-transformed values.

### Acute behaviour in pigs

Under brief anaesthesia (0.14 mg/kg intramuscular Zoletil mixture, as above), a long-line catheter (Careflow central venous catheter, Merit Medical Systems, South Jordan, UT, USA) was placed in one or both ear veins. Meloxicam (0.4 mg/kg) and amoxicillin (15 mg/kg) were given intramuscularly to minimise postoperative pain and infection, and meloxicam (0.4 mg/kg, perorally, daily) was continued for at least 3 days. Treatment was prolonged in some cases due to inflammation. Behaviour experiments began 5–10 days after catheter placement.

Animal behaviour was studied in two independent cohorts of pigs. The first group of pigs (n = 11) received DMT, psilocybin, LSD, and lisuride and the second group (n = 8) received 2-Br-LSD alone, 2-Br-LSD after volinanserin pretreatment, and lisuride after volinanserin pretreatment. Drugs were given in a randomised cross-over design, meaning that all pigs received all drugs in a randomised order with a 3–4-day washout between drug sessions to limit carry-over and allow any tolerance to subside. Not all pigs completed all drug sessions because of failure of the ear-vein catheter. Preceding all drug sessions, saline was administered alone to establish baseline behaviour. Drug doses were chosen on the basis of IC_50_ values estimated in the PET occupancy study and corresponded to approximately 60–70% 5-HT_2A_ receptor occupancy: DMT 400 µg/kg, psilocybin 30 µg/kg, LSD 0.6 µg/kg, lisuride 5.8 µg/kg, and 2-Br-LSD 11.7 µg/kg. Intravenous moclobemide (0.85 mg/kg) was given 30 min before DMT. To determine the effect of moclobemide, a [^11^C]Cimbi-36 baseline and block scans were performed in a single pig that received 0.85 mg/kg moclobemide followed 30 min later by 400 µg/kg DMT. This setup yielded a 5-HT_2A_ receptor occupancy of 42%, close to what was seen without moclobemide. Moreover, the mean plasma concentration and plasma half-life were comparable with and without moclobemide, indicating that MAO inhibition did not appreciably alter the metabolism of DMT. We therefore assumed that the measured occupancy was probably underestimated. To estimate peak occupancy at this dose, we instead used the peak plasma concentration from this pig and the IC_50_ derived from peak DMT plasma concentrations, yielding an estimated 5-HT_2A_ receptor occupancy of 73%. Volinanserin was given intravenously at 75 μg/kg, either 30 min before 2-Br-LSD or lisuride or as a co-administration. No difference was found between the two types of volinanserin administration (Supplementary Figure S8), and data were pooled for analysis.

Saline or drug was administered intravenously via the permanent ear catheter while the pigs were freely moving and awake in their home pen. The pigs were video-recorded for 30–60 min (baseline) or 60–120 min (drug) from two projections (side and above) using GoPro action cameras (HERO9 and HERO10, GoPro Inc., San Mateo, USA). Two observers blinded to the administered drug scored the occurrence of headshakes, scratching, and rubbing events from the recordings as previously described^19^.

### Statistical analysis

Headshakes, scratching, and rubbing were analysed separately using the same statistical models. Because the two experimental cohorts received different drugs and shared no animals, they were analysed as two independent datasets. The significance threshold was set at α = 0.05 and all tests were two-sided. All statistical analyses were performed in R (v. 4.6.0) using the package LMMstar (v. 1.1.0).

#### Comparison between compounds

We used linear mixed models to assess whether the compounds induced different numbers of headshakes, scratching events, or rubbing events (denoted y_d_) over the 60 min following administration. Within each data set, the model included an adjustment for the number of behaviours during the preceding saline recording (denoted y_s_), an adjustment for the duration of the saline recording (denoted t_s_) with an interaction with y_s_ because saline recordings varied in duration (minimum 30 min), and an effect of each drug.

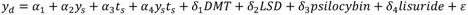

A similar model was used to test for the effects of 2-Br-LSD, 2-Br-LSD after volinanserin and lisuride after volinanserin. The residuals ε were assumed independent across pigs and to follow a compound symmetry covariance pattern within pigs. Within each data set, each compound was compared pairwise using Tukey contrasts, with p-values adjusted for multiple comparisons using a max-test procedure. To compare compounds across the two pig cohorts, the estimated mean for each drug was obtained as a linear contrast of the model coefficients and the difference between each estimate was tested using a Wald test.

Due to the randomised cross-over design, we tested whether the results were sensitive to carry-over or tolerance effects by adjusting the models for experimental day and/or compound administered in previous session, which were assumed to have a linear effect.

#### Comparison between drug and saline

We used paired t-tests to assess whether each drug induced a different number of headshakes, scratching or rubbing behaviour compared with the preceding saline recording. Each drug was tested separately. P-values were adjusted for multiple comparisons using the Benjamini-Hochberg procedure.

#### Temporal evolution of behaviour

To assess how drug effects varied over the 60 min observation period, we summed behaviour events into four consecutive 15 min intervals: 0–15, 15–30, 30–45, and 45–60 min following administration. A mean 15-min rate was calculated for each saline session and used for comparison with the drug effects; this was possible because saline behaviours were constant within the observation period. The drug effects were compared to saline effects from the preceding saline session in the same pig using paired t-tests. P-values were adjusted for multiple comparisons using the Benjamini-Hochberg procedure.

## Code availability

The code used for analyses is available via GitHub at https://github.com/camadsen/Pig_5HT2AR_Occupancy_Behaviour.

## Data availability

Data can be made available upon reasonable request.

## Author contributions

C.A.M. designed the study, collected data, performed data analysis, created visualisations, and wrote the manuscript. M.N. and E.K. assisted in collection of PET receptor occupancy and behaviour data. J.F.M.W. and C.D. collected and analysed data for in vitro 5-HT_2A_ receptor radioligand binding. V.D., V.C., and D.K. assisted in collecting and quantifying behaviour data. V.S. and M.Z. performed radiochemistry for the PET scans under supervision from M.H. D.L. and F.H. conducted quantification of DMT, LSD, ketanserin, and 2-Br-LSD plasma drug levels and provided assistance with pharmacokinetic analysis under supervision from M.L. C.B. and J.J.W. conducted quantification of volinanserin and lisuride plasma drug levels. S.S.J. conducted quantification of psilocin and LSD plasma drug levels. B.O. supervised the statistical analysis. H.D.G. designed the study and supervised the project implementation. G.M.K. designed the study, supervised the project implementation, provided resources, and contributed to manuscript writing and revision. All authors reviewed and approved the final version of the manuscript.

### Acknowledgements

We sincerely thank Dr. Cassis Varlow and Dr. Arafat Nasser for their assistance during PET scans, the animal caretakers and veterinarians at the Animal Core Facility, University of Copenhagen, for their assistance in preparing the animals for experiments, and Dr. Edis Tireli for valuable discussions on the analysis of the behavioural data and for reviewing analysis code.

