## Supplementary Material for "A Translational Porcine Model Linking Brain 5-HT_2A_ Receptor Occupancy to Psychedelic-like Behaviour"

**Supplementary figures**

- S1: 5-$HT_{2A}$ receptor occupancy from DMT bolus scans.
- S2: 5-$HT_{2A}$ receptor occupancy for DMT, psilocybin, LSD, lisuride, and 2-Br-LSD.
- S3: 5-$HT_{2A}$ receptor occupancy for volinanserin and ketanserin.
- S4: Change in mean arterial pressure (MAP) after drug administration of DMT, psilocybin, LSD, lisuride, and 2-Br-LSD.
- S5: Cumulative scores of rubbing during the 60 min following drug administration.
- S6: Agreement between in vitro $K_i$ and in vivo $IC_{50}$ values for 5-$HT_{2A}$ receptor binding
- S7: Cumulative scores of headshakes and scratching during 30 min post-drug administration of saline, moclobemide and DMT
- S8: Cumulative scores of headshakes and scratching during 30 min post-drug administration of saline, volinanserin, lisuride and 2-Br-LSD
- S9: Plasma concentrations of DMT, psilocin, LSD, lisuride and 2-Br-LSD normalized to peak plasma concentration

**Supplementary tables**

- S1: Animal and scan information for the PET experiments.
- S2: Pharmacokinetic and receptor occupancy measurements across animals.
- S3: $K_i$ and $IC_{50}$ values for DMT, psilocin, LSD, lisuride and 2-Br-LSD for in vitro binding to pig prefrontal cortex.
- S4: Estimated occupancies at the behavioural doses.

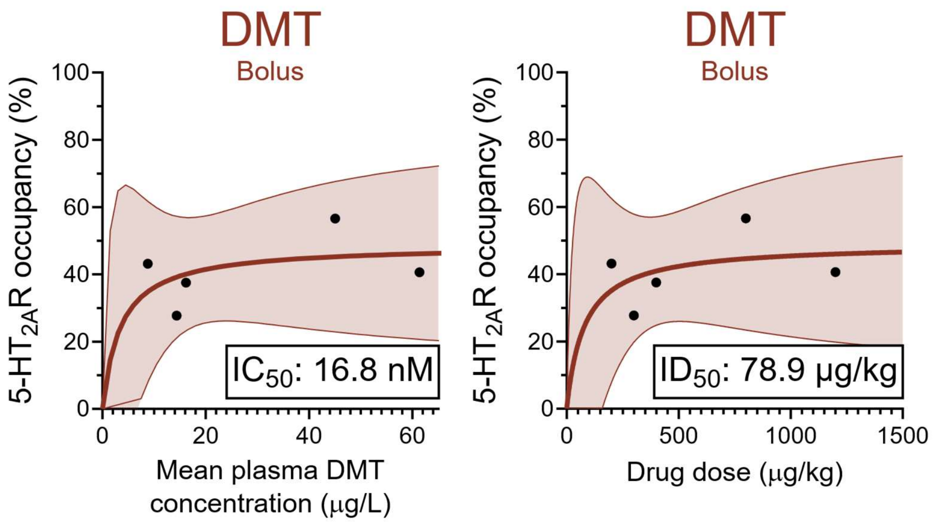


**Supplementary Figure S1. 5-$HT_{2A}$ receptor occupancy from DMT bolus scans.** Relationship between mean plasma drug concentrations over the 90 min [$^{11}$C]Cimbi-36 PET scan or drug dose and cerebral 5-$HT_{2A}$ receptor occupancy. Solid lines represent model fits, and the 95% confidence intervals are shown in coloured shading. DMT did not achieve occupancies >57%, presumably due to its short half-life.

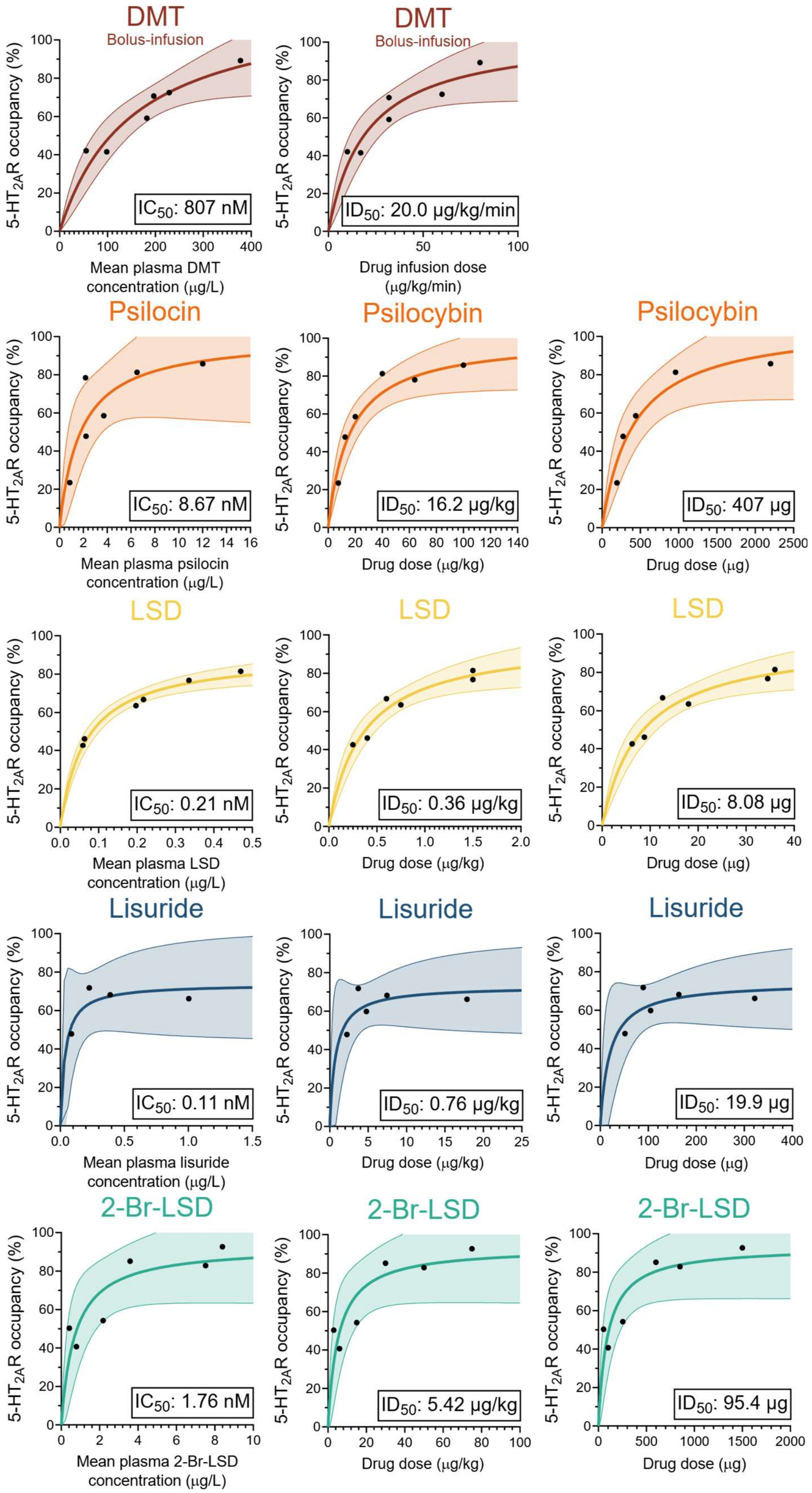


**Supplementary Figure S2. 5-$HT_{2A}$ receptor occupancy for DMT, psilocybin, LSD, lisuride, and 2-Br-LSD.** Relationship between mean plasma drug concentrations over the 90 min [$^{11}$C]Cimbi-36 PET scan or drug dose (weight-adjusted or total dose) and cerebral 5-$HT_{2A}$ receptor occupancy. Solid lines represent model fits, and the 95% confidence intervals are shown in coloured shading.

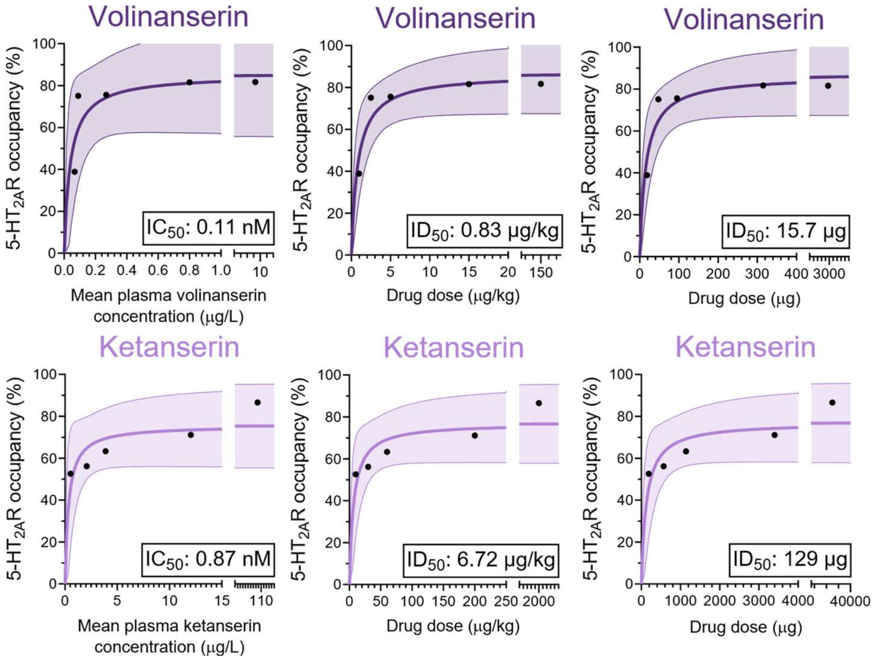


**Supplementary Figure S3. 5-HT$_{2A}$ receptor occupancy for volinanserin and ketanserin.** Relationship between mean plasma drug concentrations over the 90 min [$^{11}$C]Cimbi-36 PET scan or drug dose (weight-adjusted or total dose) and cerebral 5-HT$_{2A}$ receptor occupancy. Solid lines represent model fits, and the 95% confidence intervals are shown in coloured shading.

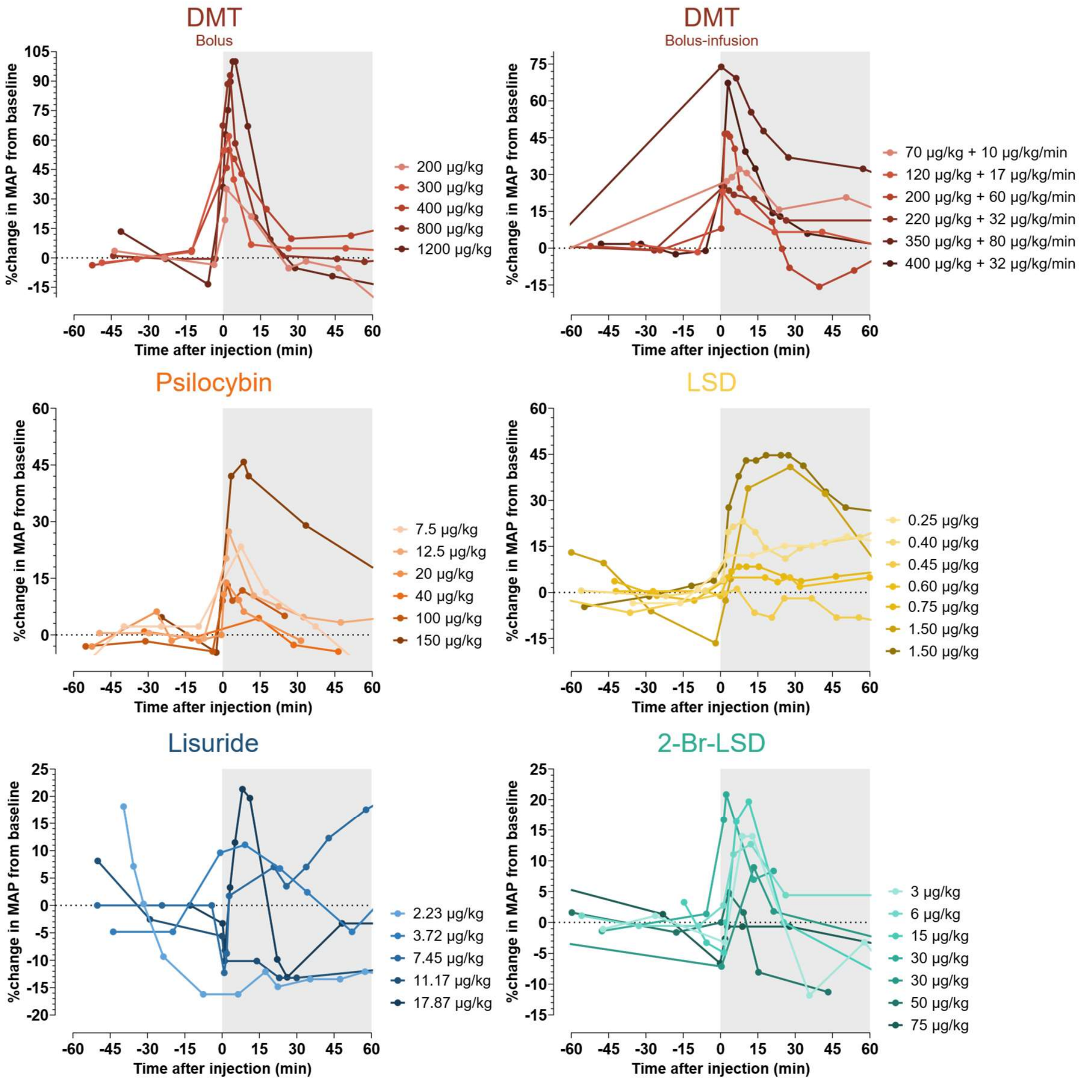


**Supplementary Figure S4. Change in mean arterial pressure (MAP) after drug administration of DMT, psilocybin, LSD, lisuride, and 2-Br-LSD.** Data are presented as the percentage change in MAP relative to the time of drug administration (min). Change in MAP is normalised to baseline for each pig; baseline was calculated as the mean of all pre-injection measurements within 60 min before drug administration.

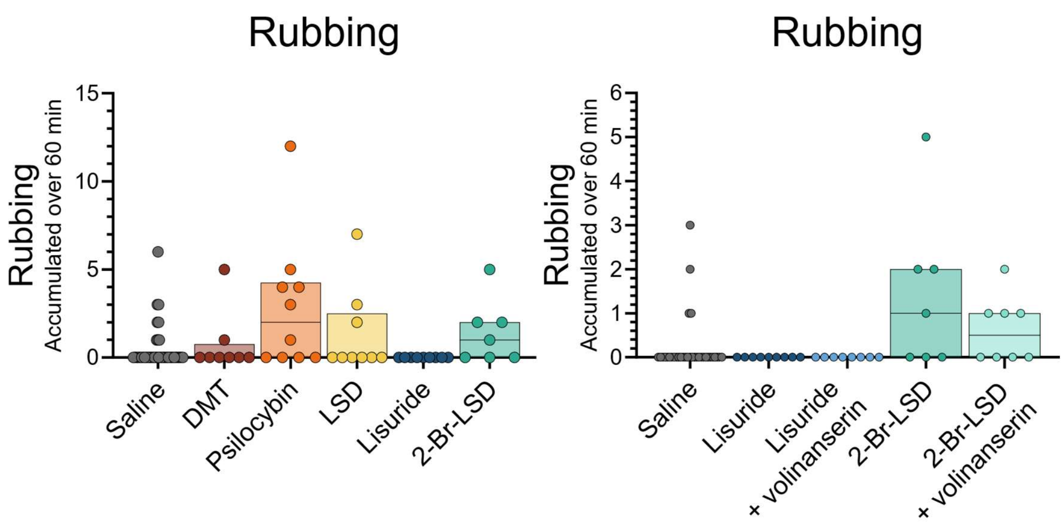


**Supplementary Figure S5. Cumulative scores of rubbing during the 60 min following drug administration.** Data are presented as median with 25th–75th percentiles.

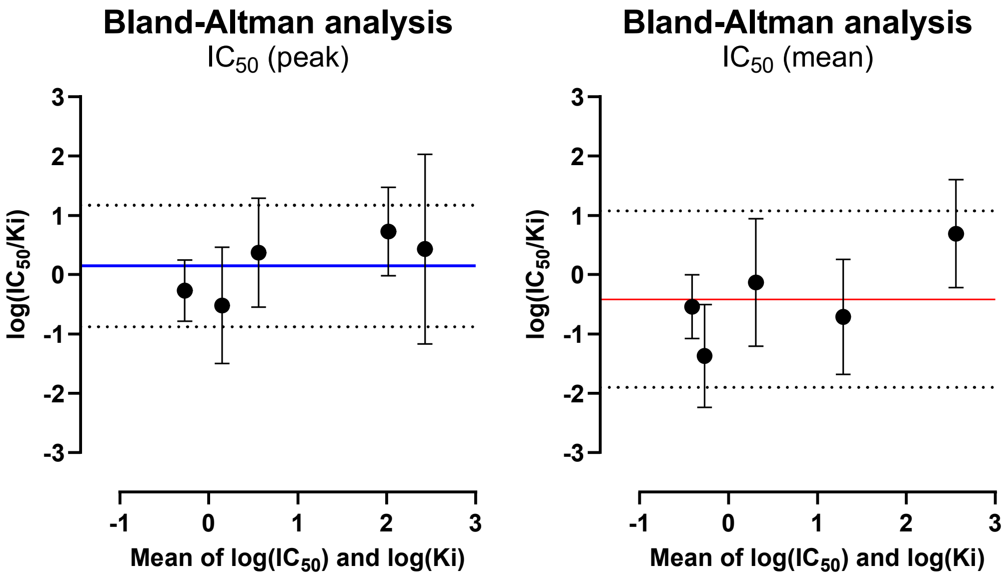


**Supplementary Figure S6. Agreement between in vitro $K_i$ and in vivo $IC_{50}$ values for 5-$HT_{2A}$ receptor binding**. Bland–Altman plots comparing in vitro $K_i$ values from [$^3$H]Cimbi-36 competition binding with in vivo $IC_{50}$ values from [$^{11}$C]Cimbi-36 PET. The $IC_{50}$ values were computed from peak plasma concentrations at scan start and mean plasma concentrations during the 90 min scan. Each data point represents one compound. The y-axis shows the difference between log $IC_{50}$ and log $K_i$, i.e., log($IC_{50}$/$K_i$), where 0 indicates perfect agreement. The x-axis shows the mean of log $IC_{50}$ and log $K_i$. Coloured lines show the bias (mean difference), and dotted lines the 95% limits of agreement. Error bars represent 95% CIs of each difference.

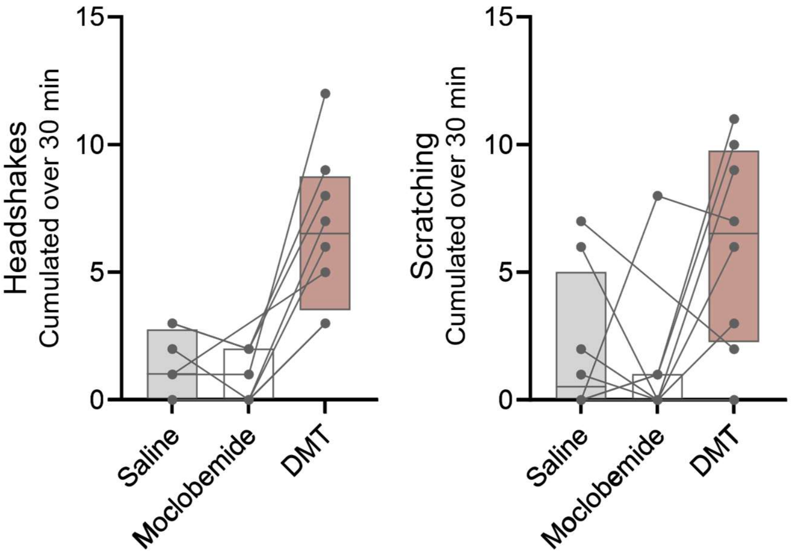


**Supplementary Figure S7. Cumulative scores of headshakes and scratching during 30 min post-drug administration of saline, moclobemide and DMT.** Data are presented as median with 25th–75th percentiles. Data from each pig is connected with a line.

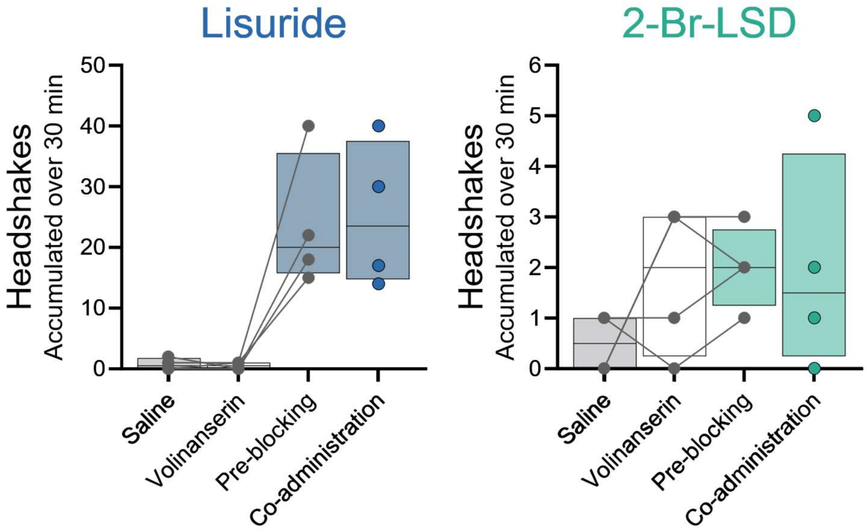


**Supplementary Figure S8. Cumulative scores of headshakes and scratching during 30 min post-drug administration of saline, volinanserin, lisuride and 2-Br-LSD.** Data are presented as median with 25th–75th percentiles. Data from each pig are connected by a line.

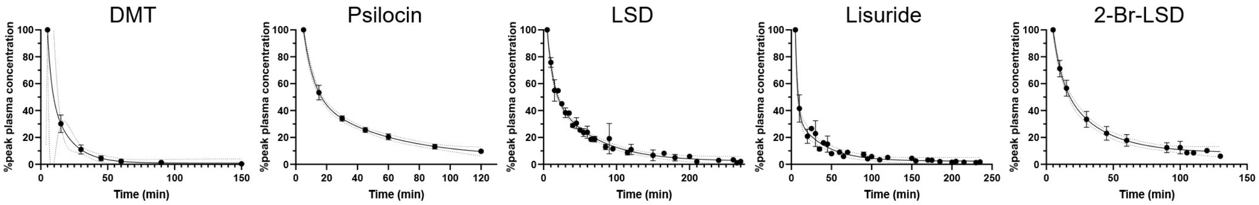


**Supplementary Figure S9. Plasma concentrations of DMT, psilocin, LSD, lisuride and 2-Br-LSD normalized to peak plasma concentration.** Peak plasma concentration for each drug was taken as the arterial plasma concentration 5 min after drug administration. Data represent mean ± s.d. A bi-exponential model is fitted to the data, plotted as the solid line. 95% confidence intervals are plotted as dotted lines.

**Supplementary Table S1. Animal and scan information for the PET experiments.**

| Pig ID | Weight | Baseline | Block scan 1 | | | Block scan 2 | | |
|---|---|---|---|---|---|---|---|---|
| | | *Injected dose (MBq)* | *Drug* | *Dose* | *Injected dose (MBq)* | *Drug* | *Dose* | *Injected dose (MBq)* |
| 17-246 | 27 kg | 574 | Psilocybin | 64 µg/kg | 399 | - | - | - |
| 18-384 | 22 kg | 486 | Lisuride | 4.77 µg/kg | 279 | - | - | - |
| 22-030 | 24 kg | 473 | Psilocybin | 40 µg/kg | 133 | - | - | - |
| 22-041 | 22 kg | 530 | Psilocybin | 20 µg/kg | 407 | - | - | - |
| 22-102 | 26 kg | 466 | Psilocybin | 7.5 µg/kg | 506 | - | - | - |
| 22-129 | 22 kg | 481 | Psilocybin | 12.5 µg/kg | 525 | Psilocybin | 100 µg/kg | 468 |
| 22-170 | 24 kg | 520 | Lisuride | 3.72 µg/kg | 566 | Follow-up | - | 513 |
| 22-196 | 23 kg | 505 | DMT | 400 µg/kg | 525 | Lisuride | 2.23 µg/kg | 484 |
| 22-207 | 24 kg | 512 | DMT | 200 µg/kg | 501 | LSD | 1.5 µg/kg | 523 |
| 22-210 | 23 kg | 542 | DMT | 300 µg/kg | 515 | LSD | 1.5 µg/kg | 569 |
| 22-252 | 24 kg | 498 | DMT | 800 µg/kg | 458 | LSD | 0.75 µg/kg | 453 |
| 22-263 | 25 kg | 471 | DMT | 1150 µg/kg | - | - | - | - |
| 22-275 | 22 kg | 474 | Lisuride | 7.45 µg/kg | 468 | Follow-up | - | 472 |
| 22-283 | 22 kg | 499 | LSD | 0.4 µg/kg | 229 | - | - | - |
| 22-326 | 22 kg | 483 | DMT | 1200 µg/kg | 484 | - | - | - |
| 23-046 | 21 kg | 481 | LSD | 0.6 µg/kg | 547 | Follow-up | - | 508 |
| 23-081 | 25 kg | 426 | LSD | 0.25 µg/kg | 503 | Follow-up | - | 455 |
| 23-228 | 18 kg | 484 | DMT | 400 µg/kg + 32 µg/kg/min | 531 | - | - | - |
| 23-264 | 19 kg | 483 | Lisuride | 11.17 µg/kg | 411 | - | - | - |
| 24-015 | 20 kg | 501 | DMT | 200 µg/kg + 60 µg/kg/min | 512 | - | - | - |
| 24-067 | 20 kg | 470 | LSD | 0.45 µg/kg | 512 | Follow-up | - | 495 |
| 24-079 | 17 kg | 454 | DMT | 220 µg/kg + 32 µg/kg/min | 460 | - | - | - |
| 24-121 | 19 kg | 531 | DMT | 70 µg/kg + 10 µg/kg/min | 379 | DMT | 120 µg/kg + 17 µg/kg/min | 469 |
| 24-274 | 17 kg | 506 | Ketanserin | 200 µg/kg | 442 | Ketanserin | 2000 µg/kg | 452 |
| 25-001 | 18 kg | 344 | 2-Br-LSD | 3 µg/kg | 524 | - | - | - |
| 25-011 | 17 kg | 499 | 2-Br-LSD | 6 µg/kg | 468 | 2-Br-LSD | 15 µg/kg | 509 |
| 25-053 | 19 kg | 501 | Ketanserin | 60 µg/kg | 384 | - | - | - |
| 25-054 | 20 kg | 508 | 2-Br-LSD | 30 µg/kg | 524 | - | - | - |
| 25-058 | 19 kg | 494 | Ketanserin | 10 µg/kg | 384 | Ketanserin | 30 µg/kg | 381 |
| 25-076 | 21 kg | 475 | Volinanserin | 15 µg/kg | 498 | Volinanserin | 150 µg/kg | 503 |
| 25-126 | 20 kg | 421 | DMT | 350 µg/kg + 80 µg/kg/min | 366 | Follow-up | - | 400 |
| 25-154 | 17 kg | 470 | 2-Br-LSD | 50 µg/kg | 423 | - | - | - |
| 25-162 | 19 kg | 429 | Volinanserin | 1 µg/kg | 392 | Volinanserin | 5 µg/kg | 418 |
| 25-164 | 20 kg | 384 | 2-Br-LSD | 30 µg/kg | 425 | 2-Br-LSD | 75 µg/kg | 377 |
| 25-185 | 18 kg | 392 | Lisuride | 17.87 µg/kg | 396 | Follow-up | - | 363 |
| 25-210 | 19 kg | 398 | Psilocybin | 150 µg/kg | 412 | Volinanserin | 2.5 µg/kg | 333 |

**Supplementary Table S2. Pharmacokinetic and receptor occupancy measurements across animals.**

| | Drug dose | Peak plasma concentration | Mean plasma concentration | Plasma half-life | Occupancy Lassen plot | Occupancy LEO |
|---|---|---|---|---|---|---|
| **DMT** (bolus) | 400 µg/kg | 407.8 µg/L | 16.2 µg/L | 17.9 min | 37.5 % | 36.0 % |
| | 200 µg/kg | 220.9 µg/L | 8.79 µg/L | 11.7 min | 43.2 % | 39.9 % |
| | 300 µg/kg | 363.4 µg/L | 14.4 µg/L | 13.6 min | 27.7 % | 22.8 % |
| | 800 µg/kg | 1019 µg/L | 45.0 µg/L | 21.7 min | 56.6 % | 56.7 % |
| | 1200 µg/kg | 1052 µg/L | 61.3 µg/L | 20.6 min | 40.6 % | 33.0 % |
| **DMT** (bolus-infusion) | 32 µg/kg/min | 285 µg/L | 182 µg/L | *NA* | 59.2 % | 47.0 % |
| | 60 µg/kg/min | 204 µg/L | 229 µg/L | *NA* | 72.5 % | 66.2 % |
| | 32 µg/kg/min | 138 µg/L | 197 µg/L | *NA* | 70.7 % | 32.5 % |
| | 10 µg/kg/min | 56.4 µg/L | 55.2 µg/L | *NA* | 42.1 % | 39.6 % |
| | 17 µg/kg/min | 88.0 µg/L | 98.3 µg/L | *NA* | 41.6 % | 35.7 % |
| | 80 µg/kg/min | 332 µg/L | 378 µg/L | *NA* | 89.2 % | 91.3 % |
| **Psilocybin** | 64 µg/kg | *NA* | *NA* | *NA* | 78.0 % | 73.1 % |
| | 40 µg/kg | 126 µg/L | 6.50 µg/L | 54.2 min | 81.3 % | 83.2 % |
| | 20 µg/kg | 63.7 µg/L | 3.71 µg/L | 45.6 min | 58.5 % | 55.7 % |
| | 7.5 µg/kg | 24.8 µg/L | 0.87 µg/L | 15.3 min | 23.5 % | 14.4 % |
| | 12.5 µg/kg | 39.5 µg/L | 2.22 µg/L | 50.0 min | 47.8 % | 47.4 % |
| | 100 µg/kg | 185 µg/L | 12.0 µg/L | 48.9 min | 85.8 % | 87.6 % |
| | 150 µg/kg | 62.1 µg/L | 2.18 µg/L | 66.9 min | 78.4 % | 74.9 % |
| **LSD** | 1.5 µg/kg | 1.40 µg/L | 0.47 µg/L | 40.5 min | 81.5 % | 76.3 % |
| | 1.5 µg/kg | 0.81 µg/L | 0.33 µg/L | 37.1 min | 76.7 % | 73.3 % |
| | 0.75 µg/kg | 0.42 µg/L | 0.20 µg/L | 41.0 min | 63.5 % | 61.7 % |
| | 0.4 µg/kg | 0.14 µg/L | 0.06 µg/L | 40.0 min | 46.2 % | 41.4 % |
| | 0.6 µg/kg | 0.37 µg/L | 0.22 µg/L | 75.1 min | 66.8 % | 67.0 % |
| | 0.25 µg/kg | 0.11 µg/L | 0.06 µg/L | 53.0 min | 42.7 % | 39.8 % |
| | 0.45 µg/kg | 0.22 µg/L | 0.16 µg/L | 53.6 min | *NA* | *NA* |
| **Lisuride** | 4.77 µg/kg | *NA* | *NA* | *NA* | 59.8 % | 56.3 % |
| | 3.72 µg/kg | 1.43 µg/L | 0.23 µg/L | 74.8 min | 71.8 % | 73.8 % |
| | 2.23 µg/kg | 0.71 µg/L | 0.09 µg/L | 51.5 min | 47.8 % | 36.0 % |
| | 7.45 µg/kg | 1.51 µg/L | 0.39 µg/L | 61.3 min | 68.1 % | 57.9 % |
| | 11.17 µg/kg | 1.11 µg/L | 0.42 µg/L | 40.7 min | *NA* | *NA* |
| | 17.87 µg/kg | 6.03 µg/L | 1.00 µg/L | 59.0 min | 66.2 % | 58.7 % |
| **2-Br-LSD** | 3 µg/kg | 1.30 µg/L | 0.42 µg/L | 87.6 min | 50.3 % | 49.4 % |
| | 6 µg/kg | 2.47 µg/L | 0.79 µg/L | 66.5 min | 40.7 % | 28.3 % |
| | 15 µg/kg | 6.16 µg/L | 2.18 µg/L | 50.6 min | 54.3 % | 42.1 % |
| | 30 µg/kg | 11.2 µg/L | 2.65 µg/L | 67.0 min | *NA* | *NA* |
| | 50 µg/kg | 25.0 µg/L | 7.52 µg/L | 45.2 min | 82.8 % | 75.1 % |
| | 30 µg/kg | 13.9 µg/L | 3.58 µg/L | 51.7 min | 85.1 % | 80.4 % |
| | 75 µg/kg | 33.2 µg/L | 8.39 µg/L | 49.2 min | 92.6 % | 89.5 % |
| **Volinanserin** | 15 µg/kg | 4.56 µg/L | 0.80 µg/L | 56.5 min | 81.6 % | 83.5 % |
| | 150 µg/kg | 47.2 µg/L | 9.63 µg/L | 59.2 min | 81.7 % | 81.9 % |
| | 1 µg/kg | 0.36 µg/L | 0.07 µg/L | 46.2 min | 38.9 % | 38.3 % |
| | 5 µg/kg | 1.30 µg/L | 0.27 µg/L | 55.4 min | 75.6 % | 75.3 % |
| | 2.5 µg/kg | 0.29 µg/L | 0.09 µg/L | 49.3 min | 75.1 % | 75.8 % |
| **Ketanserin** | 200 µg/kg | 24.4 µg/L | 12.0 µg/L | 39.1 min | 71.1 % | 62.9 % |
| | 2000 µg/kg | 208 µg/L | 109 µg/L | 49.7 min | 86.6 % | 71.5 % |
| | 60 µg/kg | 10.8 µg/L | 3.88 µg/L | 45.5 min | 63.3 % | 57.9 % |
| | 10 µg/kg | 2.61 µg/L | 0.54 µg/L | 23.4 min | 52.6 % | 54.7 % |
| | 30 µg/kg | 4.96 µg/L | 2.07 µg/L | 33.2 min | 56.2 % | 55.6 % |

**Supplementary Table S3. $K_i$ and $IC_{50}$ values for DMT, psilocin, LSD, lisuride and 2-Br-LSD for in vitro binding to pig prefrontal cortex.**

| | DMT | Psilocin | LSD | Lisuride | 2-Br-LSD |
|---|---|---|---|---|---|
| $K_i$ (mean ± SD) | 489.0 ± 614.7 nM | 67.1 ± 62.0 nM | 1.5 ± 2.1 nM | 3.3 ± 2.5 nM | 3.6 ± 2.8 nM |
| $IC_{50}$ (mean ± SD) | 2370 ± 2979 nM | 325 ± 300 nM | 7.2 ± 10.2 nM | 15.8 ± 11.9 nM | 17.4 ± 13.7 nM |

**Supplementary Table S4. Estimated occupancies at the behavioural doses.**

| Drug | Behaviour dose | Estimated 5-$HT_{2A}$ receptor occupancy |
|---|---|---|
| DMT | 400 μg/kg | 73.1 % |
| Psilocybin | 30 μg/kg | 65.5 % |
| LSD | 0.6 μg/kg | 61.3 % |
| Lisuride | 5.8 μg/kg | 64.5 % |
| 2-Br-LSD | 11.7 μg/kg | 63.8 % |
| Volinanserin | 75 μg/kg | 85.4% |
